# CDK/mTOR–dependent phosphorylation of UBE2H restrains its charging with ubiquitin and regulates CTLH-dependent degradation

**DOI:** 10.64898/2026.03.07.710281

**Authors:** Yingqian Chen, Valentina Rossio, Joao A. Paulo, Menuka Karki, Sandhya Manohar, Noelle Ozimek, Laura Frizzi, Steven P. Gygi, Randall W. King

## Abstract

The C-terminal to LisH (CTLH) complex is a modular multi-subunit E3 ligase with diverse biological functions, yet how its overall ubiquitylation activity is tuned remains unclear. Here, we identify CDK- and mTOR-dependent phosphorylation of the cognate E2 enzyme UBE2H as a key regulator of CTLH E3 catalytic capacity. Phosphorylation of two N-terminal serine residues (S3/S5) reduces UBE2H charging with ubiquitin, thereby limiting the pool of active E2 available to CTLH. Mitotic CDK activity inactivates UBE2H during mitosis, whereas mTOR restrains UBE2H charging in interphase to couple CTLH-dependent ubiquitylation to nutrient status. Preventing this phosphorylation maintains UBE2H charging, enhances CTLH-mediated substrate degradation, promotes CTLH subunit turnover, and causes proliferation and mitotic defects. Using hyperactive UBE2H, we identify two additional CTLH substrates, the mitotic kinase NEK9 and Angio-associated migratory cell protein (AAMP) and define a DR-like C-degron recognized by the CTLH subunit MKLN1. These findings reveal how regulation of an E2 enzyme by cell cycle and nutrient signaling pathways dynamically shape CTLH activity.

## Introduction

The ubiquitin system plays a central role in regulating protein homeostasis and signaling and is dynamically modulated depending on physiological context. Ubiquitylation is mediated by an enzymatic cascade in which E1s activate ubiquitin, E2s conjugate ubiquitin, and E3s recruit substrates and activate ubiquitin transfer^1,2^. Ubiquitin conjugation to substrates is typically regulated at the level of the E3 or the substrate^3–5^. For example, the activity of the Anaphase-Promoting Complex/Cyclosome (APC/C) is regulated by cell-cycle state through modulation of expression and phosphorylation of its activator proteins^6–8^. In addition, phosphorylation of Skp1–Cullin–F-box (SCF) substrates can generate phosphodegrons that promote their recognition and ubiquitylation^8^. In contrast, how E2 catalytic activity is regulated *in vivo* remains less extensively characterized. Notably, prior work has demonstrated that phosphorylation of E2 enzymes can impact their ability to interact with E1 or E3s, but the physiological significance of this regulation has not been fully explored^9^.

In living cells, ubiquitin pathway activity is often inferred through measurements of substrate ubiquitylation or degradation, but these readings are often indirect due to substrate accessibility and localization. Conversely, thioester intermediates in the ubiquitylation process (E1∼Ub, E2∼Ub, and HECT/RBR type E3∼Ub) represent the catalytically occupied states necessary for the cascade reaction, and their steady-state level is determined by the balance between charging and discharging^10,11^. Many studies have used the kinetics or steady-state levels of E2-Ub thioester formation to assess E2 activity, mutation effects, and regulatory mechanisms^12,13^. Therefore, quantitatively assessing the intracellular ubiquitin-loaded enzyme pool can provide a complementary biochemical readout of pathway readiness. However, quantitative analysis of ubiquitin charging states during the cell cycle remains limited. We therefore performed systematic, quantitative profiling of ubiquitin charging to identify cell-cycle-regulated components.

The CTLH complex is a conserved multi-subunit E3 ligase that works in partnership with its cognate E2 enzyme UBE2H. CTLH/UBE2H-dependent degradation regulates diverse biological processes, including cell proliferation, metabolism, migration and development^14–17^. In *Drosophila*, increased expression of the UBE2H ortholog Marie Kondo promotes clearance of maternal RNA-binding proteins during the maternal-to-zygotic transition^18,19^. In mammalian erythropoiesis, stage-specific remodeling of CTLH complexes and elevated UBE2H levels promote terminal maturation by remodeling the proteome^20^. CTLH activity and substrate specificity are also modulated by cell-cycle and metabolic signals. For example, the CTLH complex restrains G1 progression and limits entry into quiescence, suggesting that its activity is dynamically regulated across cell cycle states^21^. In parallel, CTLH activity is linked to nutrient status, as the complex modulates glycolytic pathways through ubiquitylation of PKM2 and LDHA^22^. Furthermore, mechanistic target of rapamycin (mTOR) inhibition enhances CTLH-dependent turnover, as reflected by loss of both MKLN1, a CTLH component that might function as both a substrate receptor and a substrate^19,23,24^, and metabolic enzymes like 3-hydroxy-3-methylglutaryl (HMG)-coenzyme A (CoA) synthase 1 (HMGCS1)^25–28^. However, the mechanisms by which mTOR signaling regulates CTLH activity remain unclear. Consistent with a conserved role in nutrient-responsive signaling, the yeast ortholog of CTLH, the glucose-induced degradation-deficient (GID) complex, was originally identified as a nutrient-responsive ubiquitin ligase that targets gluconeogenic enzymes via the substrate receptor Gid4^29,30^. Interestingly, both UBC8 and UBE2H are positively regulated by phosphorylation of their C-termini, which promotes association with the CTLH complex. However, this phosphorylation seems to be constitutive and does not explain regulation of CTLH activity^31^. Together, these observations indicate that CTLH integrates cues from cell cycle, metabolic and developmental conditions, yet the specific mechanisms that modulate CTLH activity as well how its E2 partnership is regulated remain incompletely understood.

Here, we report a phosphorylation-dependent mechanism that links cell-cycle and nutrient signaling to CTLH E3 ligase activity. We show that Cyclin-dependent kinase (CDK) and mTOR-dependent phosphorylation of the cognate E2, UBE2H, restrains CTLH-dependent degradation by limiting E2 charging. This establishes UBE2H phosphorylation as a regulatory switch that tunes CTLH activity in a context-dependent manner, providing an example of context-dependent regulation of E2 enzymes. Using a hyperactive non-phosphorylatable UBE2H mutant, we further expand the CTLH substrate repertoire and define a new degron-recognition paradigm.

## Results

### UBE2H charging is reduced during mitosis

To systematically detect quantitative differences in ubiquitin charging of E1, E2 and E3 enzymes between interphase and mitosis, we designed an *in vitro* screening assay using *Xenopus* egg extract. We added HA-tagged ubiquitin to interphase or mitotic extract and subsequently immunodepleted the extract with anti-HA or control beads. Then, we used TMT-based proteomics profiling to compare the amount of each protein remaining in the supernatant. We reasoned that a higher degree of ubiquitin-thioester formation would result in a greater depletion by anti-HA antibody (Fig. 1a). This analysis showed substantial depletion of E1s, E2s, and some HECT-based E3s, which are known to form ubiquitin thioesters (Extended Data Fig. 1a, Supplementary Table 1). Most E2 enzymes were depleted to a similar extent in interphase and mitotic extracts. In addition, several E2 enzymes that are not expected to be ubiquitin-charged were not immunodepleted (e.g. UBE2M, UBE2V1 and UBEV2). Notably, depletion of UBE2H was markedly reduced during mitosis, suggesting it may be charged less efficiently in mitosis (Fig. 1b). To determine whether these differences in depletion arose from differences in ubiquitin thioester formation, we added radiolabeled E2s expressed in reticulocyte lysate and performed non-reducing gels and autoradiography. Whereas UBE2C was charged during both interphase and mitosis, UBE2H was charged during interphase, but was predominantly uncharged during mitosis, consistent with the mass spectrometry-based results (Fig. 1c).

**Fig. 1.**
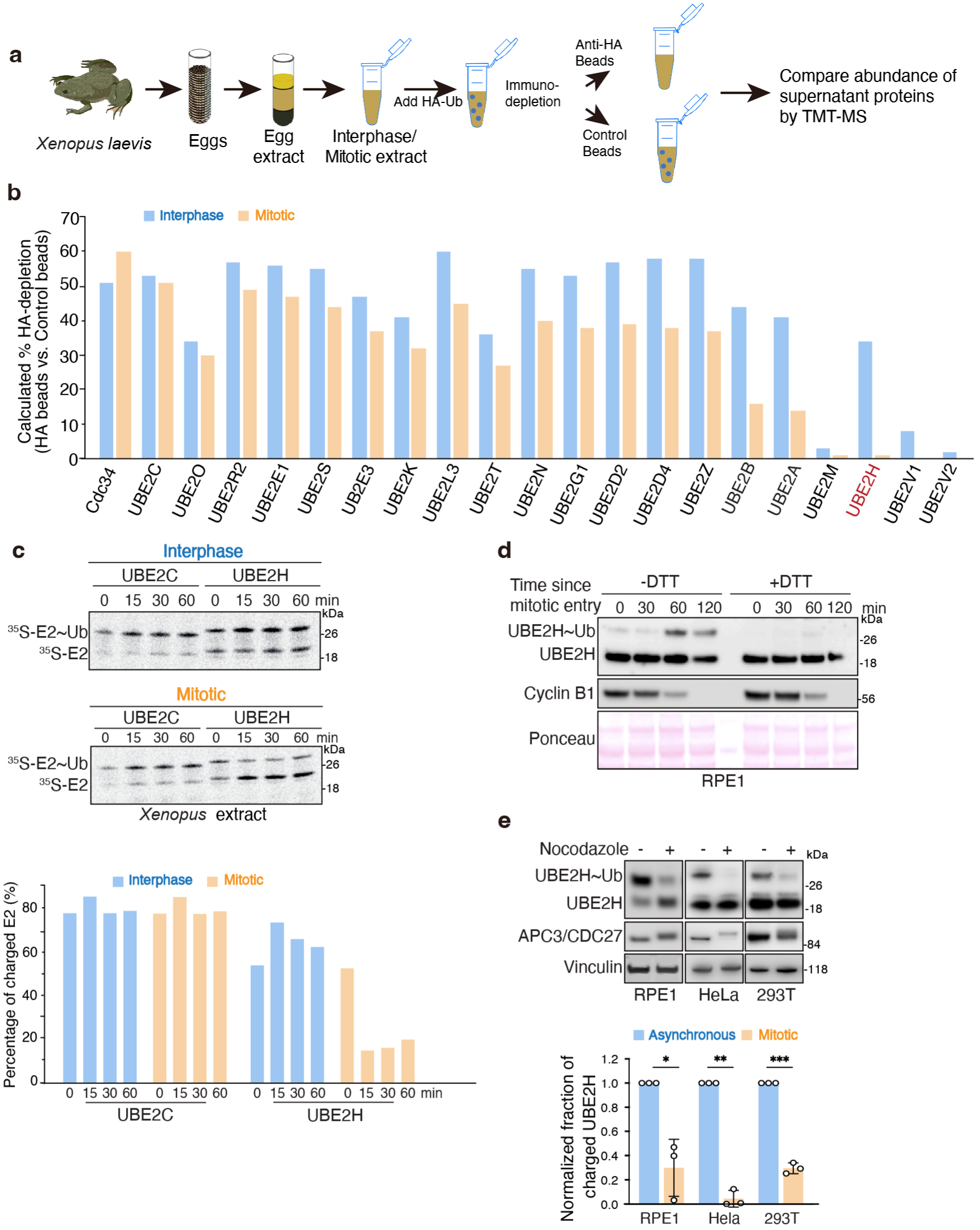
UBE2H charging is reduced during mitosis. **a.** Schematic of the proteome-wide screen for ubiquitin-bound proteins in Xenopus egg extract. HA-tagged ubiquitin was added to interphase or mitotic extracts, which were subsequently immunodepleted with anti-HA beads or control antibody beads. TMT-based proteomic profiling was used to compare the abundance of each protein remaining in the supernatant. **b.** Percent depletion of different E2 enzymes in interphase and mitotic Xenopus egg extract. The E2 showing the most pronounced reduction in depletion during mitosis, UBE2H, is highlighted in red. **c.** Levels of ubiquitin-charged recombinant radiolabeled E2s (UBE2C∼Ub or UBE2H∼Ub) were examined by autoradiography. Extracts were in interphase or pre-arrested mitosis, E2s were added and incubated for the indicated times. Reactions were quenched with non-reducing sample buffer and levels of E2∼Ub determined (top). Quantification of the percentage of ubiquitin-charged E2s is shown (bottom). **d.** Levels of ubiquitin-charged endogenous UBE2H (UBE2H ∼Ub) in hTERT-RPE1 cells were examined by anti-UBE2H immunoblotting. Cells were arrested in G2 with RO3306 (7.5 µM) for 18h, then released for 30 minutes to allow for mitotic entry. Mitotic cells were collected by mitotic shake-off, re-plated and collected at the indicated time points. Cell lysates were analyzed under non-reducing conditions to examine UBE2H∼Ub or under reducing conditions to examine total UBE2H level. **e.** Levels of ubiquitin-charged endogenous UBE2H in hTERT-RPE1, HeLa and HEK293T cells. Cells were treated with nocodazole (600 nM) for 18h, and then mitotic cells were collected by shake-off. Cell lysates were analyzed under non-reducing conditions to examine levels of UBE2H∼Ub (top). Quantification represents mean ± s.d. from three independent biological replicates. Statistical significance of differences between groups was determined with two-tailed unpaired Student’s t-test. *** = p < 0.001. ** = p < 0.01. * = p < 0.05. ns = not statistically significant.

To investigate whether UBE2H charging varies between interphase and mitosis in human RPE1 cells, we used RO3306, a CDK1 inhibitor, to arrest cells at the G2/M phase boundary. Upon release from RO3306, cells entered mitosis, as indicated by the initial accumulation of Cyclin B1, followed by a decrease in Cyclin B1 levels during mitotic exit. Accordingly, UBE2H was uncharged during mitosis, and became recharged as cells exited mitosis. We further used dithiothreitol (DTT), a reducing agent, to confirm that the higher molecular weight band represents a thioester bond with ubiquitin (Fig. 1d). Additionally, a comparable difference in UBE2H charging between asynchronous and mitotically-arrested cells was observed in HeLa and HEK293T cells (Fig. 1e). We next investigated whether UBE2H charging also varies across the different stages of interphase. In both RPE1 and HeLa cells, we observed that UBE2H is specifically discharged during mitosis, while it remains substantially charged in interphase (Extended Data Fig. 1b). Taken together, these findings indicate that in both *Xenopus* extract and human cells, UBE2H is charged and active during interphase, but becomes discharged and inactive in mitosis.

### N-terminal phosphorylation inhibits UBE2H charging during mitosis

To understand the mechanism by which UBE2H becomes discharged during mitosis, we first tested whether protein phosphorylation could be involved. We therefore treated RPE1, HeLa and 293T cells with okadaic acid, a protein phosphatase inhibitor, and examined UBE2H-ubiquitin thioester formation. We found that UBE2H became discharged rapidly upon okadaic acid treatment (Extended Data Fig. 2a). Furthermore, prior work reported that UBE2H can be phosphorylated by CDK at two conserved N-terminal serine residues (S3 and S5), which may influence its ability to be charged by E1^9^. We confirmed that recombinant CDK2/Cyclin A can phosphorylate UBE2H in an *in vitro* phos-tag gel assay (Extended Data Fig. 2b). Based on these results, we hypothesized that phosphorylation of these serine residues may underlie the loss of UBE2H activity during mitosis. To address this, we generated single (S3A, S5A) and double (AA) serine-to-alanine UBE2H mutants (Fig. 2a). We expressed radiolabeled versions of these human proteins in reticulocyte lysate and added them to *Xenopus* egg extract. We found that the individual S3A and S5A single mutants were largely unable to rescue thioester bond formation of UBE2H in mitosis, whereas the double mutant (AA) remained largely charged during mitosis (Fig. 2b). We also generated a double aspartic acid mutant (DD) to mimic constitutive phosphorylation. We found that the DD mutant remained discharged in both interphase and mitosis (Fig. 2c). Altogether, these results suggest that phosphorylation of these serine residues (S3 and S5) explains the loss of the UBE2H thioester in mitotic *Xenopus* egg extracts.

**Fig. 2.**
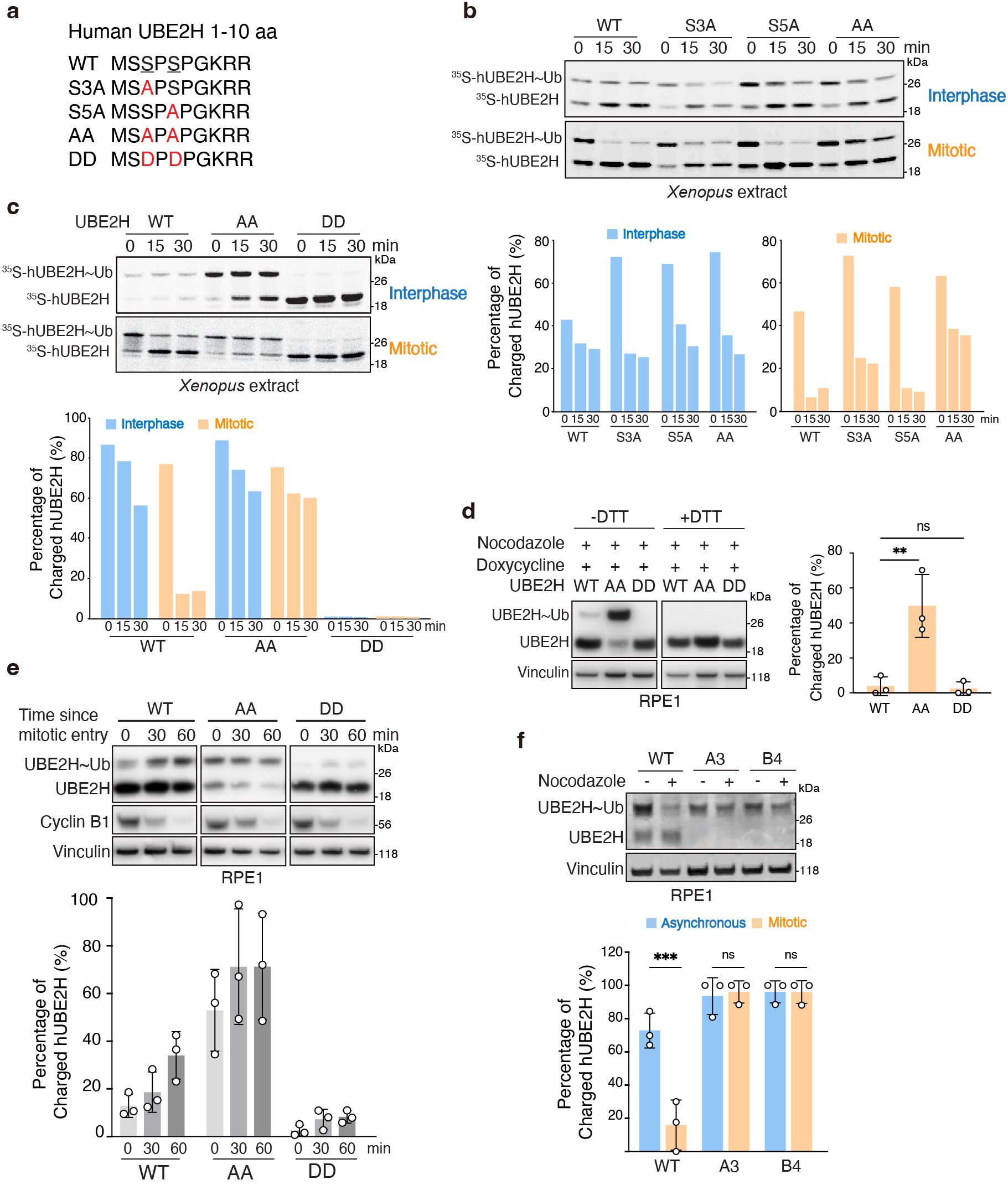
N-terminal phosphorylation inhibits UBE2H charging during mitosis. **a.** N-terminal amino acid sequence of human UBE2H (aa 1-10). Mutated residues are highlighted in red. **b.** Levels of ubiquitin-charged recombinant radiolabeled UBE2H (WT, S3A, S5A and AA) were examined by autoradiography. Xenopus egg extract was in interphase or pre-arrested mitosis, E2s were added and incubated for the indicated times and quenched with non-reducing sample buffer to examine levels of UBE2H∼Ub (top). Quantification of the percentage of ubiquitin-charged UBE2H is shown (bottom). **c.** Levels of ubiquitin-charged recombinant radiolabeled UBE2H (WT, AA and DD) were examined by autoradiography in interphase or mitotic Xenopus egg extract. **d.** Levels of doxycycline-inducible ubiquitin-charged UBE2H (WT, AA and DD) were examined by anti-UBE2H immunoblotting in stable hTERT-RPE1 cells. Cells were treated with nocodazole (600 nM) for 18h and mitotic cells were collected by shake-off. Cell lysates were analyzed under non-reducing conditions to examine UBE2H∼Ub or reducing conditions to examine total UBE2H levels. **e.** Levels of doxycycline-inducible ubiquitin-charged UBE2H (WT, AA and DD) were examined by anti-UBE2H immunoblotting in stably expressing hTERT-RPE1 cells. Cells were arrested by RO3306 (7.5 µM) for 18h, then released for the indicated times, and extracts were analyzed by immunoblotting under non-reducing conditions. Quantification represents mean± s.d. from three independent biological replicates. **f.** Two UBE2H-AA mutant clones (A3, B4) were generated using a CRISPR/-Cas9-based knock-in approach. Cells were treated with nocodazole (600 nM) for 24h to induce arrest in mitosis, collected and processed for non-reducing SOS-PAGE. Quantification represents mean ± s.d. from three independent biological replicates. Statistical significance of differences between groups was determined with one-way ANOVA analysis **(d)** or two-tailed unpaired Student’s t-test **(f).** *** = p < 0.001. ** = p < 0.01. ns = not statistically significant.

To determine whether UBE2H is regulated similarly in human cells, we used lentiviral transduction to establish stable RPE1 cells that express doxycycline-inducible WT, AA or DD constructs. In parallel, we used CRISPR-Cas9 to edit the endogenous UBE2H locus to express the AA mutation (creating two independent clones named A3 and B4) (Extended Data Fig. 2c). In the inducible cell lines, upon nocodazole-induced mitotic arrest, WT was predominantly uncharged, whereas the AA mutant remained mostly charged. As expected, the DD mutant remained uncharged in asynchronous and mitotic cells. We used reducing gels to confirm that the higher-molecular weight species of UBE2H represents a ubiquitin thioester (Fig. 2d). As we observed for the endogenous protein, exogenously expressed UBE2H-WT was discharged upon entry into mitosis induced by release from RO3306 and became recharged as cells exited mitosis. In contrast, the AA mutant remained mostly charged during these transitions, whereas the DD mutant remained mostly uncharged (Fig. 2e). In addition, we titrated the concentration of doxycycline and found that UBE2H-AA was more charged than UBE2H-WT under all concentrations in asynchronous conditions (Extended Data Fig. 2d), suggesting that interphase phosphorylation of UBE2H may also partially restrain its activity. We observed similar results in the CRISPR-edited clones, where the AA mutant clones were highly charged in both asynchronous and mitotic populations (Fig. 2f). Furthermore, whereas okadaic acid strongly suppressed ubiquitin charging of WT UBE2H, both edited clones showed less suppression of charging (Extended Data Fig. 2e). Therefore, we conclude that UBE2H phosphorylation at S3 and S5 inactivates UBE2H during mitosis and may also partially suppress UBE2H charging during interphase.

Interestingly, we observed that the protein levels of UBE2H-AA in the knock-in clones were lower than those of WT cells. To investigate whether this was due to differences in mRNA synthesis or protein turnover, we first measured *UBE2H* mRNA levels and found that expression was reduced in both A3 and B4 clones compared to WT cells (Extended Data Fig. 2f). In contrast, upon treatment with the proteasome inhibitor MG262, both WT and AA accumulated at similar rate (Extended Data Fig. 2g). Furthermore, performing a cycloheximide chase revealed that the protein turnover rates of UBE2H-WT and AA were similar (Extended Data Fig. 2h). These results indicate that UBE2H-AA expression appears to be transcriptionally repressed in the knock-in cells, suggesting the possibility that expression of the mutant protein may be deleterious to cells. Taken together, our findings suggest that phosphorylation of UBE2H may constrain its activity in interphase and strongly suppress its activity in mitosis.

### mTOR signaling restrains UBE2H charging to regulate CTLH-dependent degradation

Our previous results showed that the non-phosphorylatable UBE2H-AA mutant exhibited a higher level of ubiquitin charging than WT not only in mitosis but also in asynchronous populations, which are predominantly in interphase. This prompted us to ask whether UBE2H dephosphorylation is associated with increased CTLH-dependent turnover in interphase. Indeed, the CTLH substrate HMGCS1 was downregulated in cells overexpressing UBE2H-AA to a greater extent than cells expressing UBE2H-WT or DD (Fig. 3a). Furthermore, HMGCS1 was also downregulated in the two UBE2H-AA knock-in cell lines relative to WT counterparts (Fig. 3b). Given prior evidence linking mTOR inhibition to enhanced CTLH-associated turnover of HMGCS1, we hypothesized that mTOR signaling may restrain CTLH output in interphase by maintaining N-terminal phosphorylation of UBE2H.

**Fig. 3.**
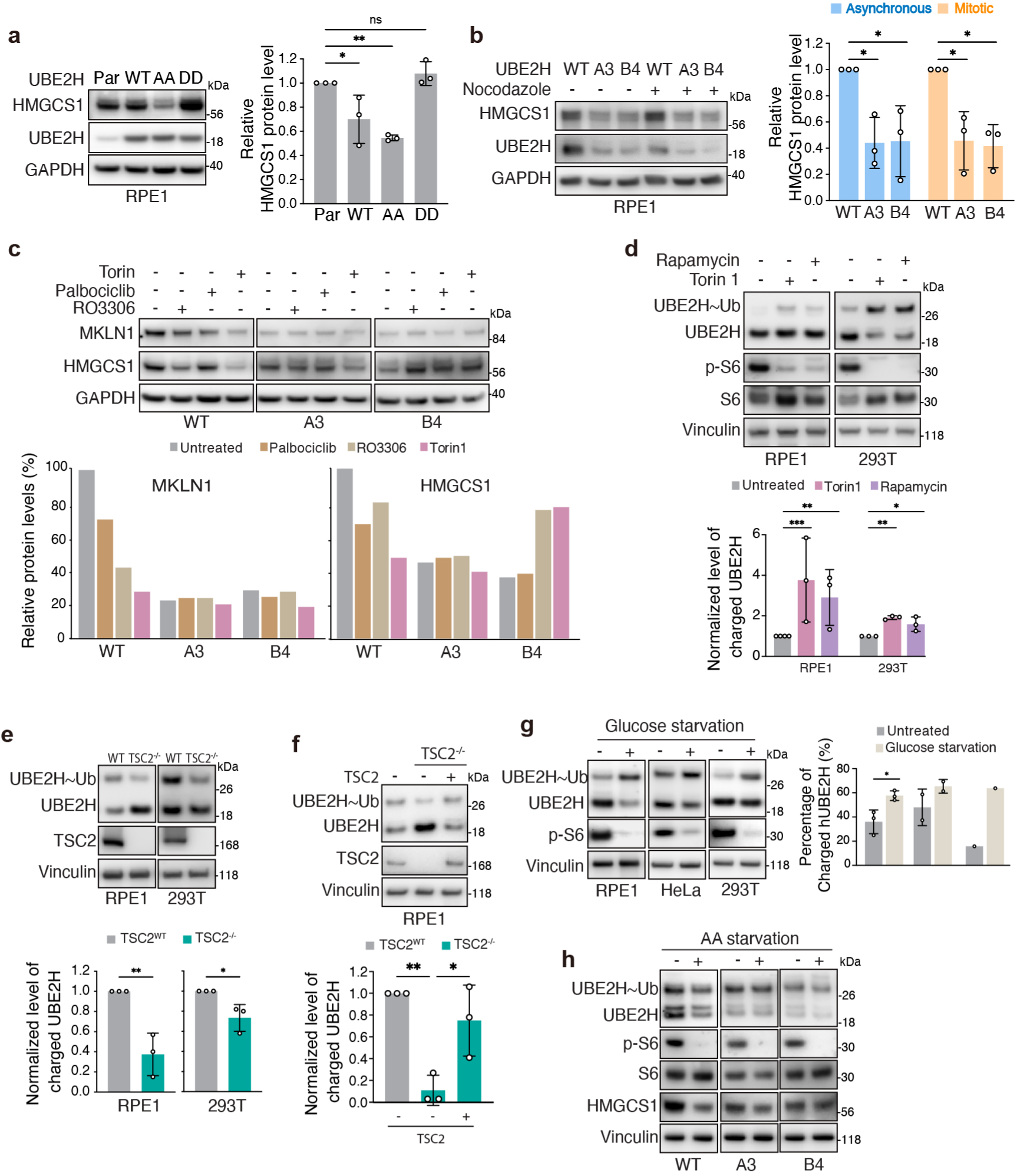
mTOR signaling restrains UBE2H charging to limit CTLH-dependent degradation. **a.** Western blots showing HMGCS1 protein level in asynchronous hTERT-RPE1 cells that stably express doxycycline-inducible UBE2H constructs (WT, AA and DD). Cells were induced with doxycycline (2.5 ng/ml) for 24 h. Quantification represents mean± s.d. from three independent biological replicates. **b.** Western blots showing HMGCS1 protein levels in asynchronous or mitotically arrested WT cells and knock-in clones. To induce mitotic arrest, cells were treated with nocodazole (600 nM) for 18h, and mitotic cells were collected by shake-off. Quantification represents mean ± s.d. from three independent biological replicates. **c.** Western blots showing MKLN1 and HMGCS1 protein levels in WT cells and knock-in clones after treatment with torin1 (250 nM), R03306 (7.5 µM) and palbociclib (1 µM) for 18h. Quantification of relative MKLN1and HMGCS1 protein levels is shown (bottom). **d.** Non-reducing SOS-PAGE analysis of UBE2H thioester-linked ubiquitin conjugates in hTERT-RPE1 cells and HEK293T cells after treatment with torin1 (250 nM) or rapamycin (80 nM) and the proteasome inhibitor MG262 for 1h. Quantification of normalized level of charged UBE2H is shown as mean± s.d. from three independent biological replicates. **e.** Non-reducing SOS-PAGE analysis of UBE2H thioester-linked ubiquitin conjugates in WT and *TSC2-^1^-* hTERT-RPE1cells and HEK293T cells. Quantification of normalized level of charged UBE2H is shown as mean± s.d. from three independent biological replicates. **f.** Non-reducing SOS-PAGE analysis of UBE2H thioester-linked ubiquitin conjugates in WT, TSC2-^1^-and *TSC2-^1^-* + TSC2 hTERT-RPE1cells. Quantification of normalized level of charged UBE2H is shown as mean± s.d. from three independent biological replicates. **g.** Non-reducing SOS-PAGE analysis of UBE2H thioester-linked ubiquitin conjugates and phospho-S6 (p-S6) in hTERT-RPE1, Hela and HEK293T cells that were incubated in glucose-depleted medium for 6h. Quantification represents mean± s.d. from the indicated independent biological replicates. **h.** Non-reducing SOS-PAGE analysis of UBE2H thioester-linked ubiquitin conjugates and phos-pho-S6 (p-S6) in WT cells and knock-in clones that were incubated in amino-acid (AA)-depleted medium for 6h. Statisti-cal significance of differences between groups was determined with one-way ANOVA analysis **(a-b,d, f)** or two-tailed unpaired Student’s t-test **(e, g).** *** = p < 0.001. ** = p < 0.01. * = p < 0.05. ns = not statistically significant.

To explore upstream kinase pathways that regulate CTLH-dependent degradation in asynchronous populations, we treated both WT and knock-in cells with inhibitors of mTOR or CDK and monitored CTLH-related readouts such as HMGCS1 and MKLN1 levels. In WT cells, the mTOR inhibitor torin1 reduced both HMGCS1 and MKLN1 levels, which is consistent with previous reports^25,27^. The CDK1 inhibitor RO3306 also downregulated HMGCS1 and MKLN1 levels like torin1, whereas the CDK4/6 inhibitor palbociclib only modestly downregulated HMGCS1 but strongly reduced the levels of MKLN1. Therefore, inhibition of either mTOR or CDK appears to be able to stimulate aspects of CTLH-dependent degradation. Interestingly, in both AA clones, inhibitor treatment did not lead to further downregulation of HMGCS1 or MKLN1 levels (Fig. 3c). These findings suggest that phosphorylation of UBE2H at S3/S5 functions downstream of CDK and mTOR signaling to regulate CTLH-dependent degradation.

Next, we tested whether kinase inhibition impacts UBE2H charging by measuring the thioester formation of UBE2H. Inhibition of mTOR with torin1 and rapamycin increased UBE2H thioester formation in both RPE1 and 293T cells. This was accompanied by a reduction in phospho-S6 levels, consistent with inhibition of mTORC1 signaling (Fig. 3d and Extended Data Fig. 3a-b). We then asked whether increased mTOR signaling is sufficient to suppress UBE2H charging. To test this, we analyzed TSC2 knockout RPE1 and 293T cells, which display constitutively activated mTOR signaling^32^. Loss of TSC2 decreased UBE2H thioester formation (Fig. 3e), and this effect was restored by stable re-expression of TSC2 (Fig. 3f). Furthermore, treating cells with mTOR inhibitors partially restored UBE2H thioester formation in the TSC2-knockout background (Extended Data Fig. 3c). We also tested how physiological inhibition of mTOR activity influenced UBE2H thioester formation. We found that starvation of glucose (Fig. 3g), amino acids (Fig. 3h) or serum (Extended Data Fig. 3d) were each sufficient to increase UBE2H thioester formation. Importantly, these effects were largely blunted in the UBE2H knock-in cells, consistent with UBE2H S3/S5 phosphorylation mediating the mTOR-dependent regulation of UBE2H charging. Taken together, our findings suggest that mTOR signaling restrains UBE2H charging, thereby regulating CTLH-associated turnover in asynchronous cell populations.

### Cells that cannot phosphorylate UBE2H show defects in proliferation and mitosis

We next asked what happens to cells if they cannot phosphorylate and inactivate UBE2H. First, we observed that both the AA inducing cells and the knock-in clones exhibited slower growth compared to WT cells (Fig. 4a-b), accompanied by an increased fraction of cells in G1 phase (Extended Data Fig. 4a) and lower colony formation ability (Extended Data Fig. 4b). Since our data suggest that UBE2H is normally inactivated during mitosis, we also investigated how expression of hyperactive, non-phosphorylatable UBE2H affected chromosome segregation, mitotic duration, and spindle checkpoint function. Time lapse imaging of AA knock-in cells demonstrated an increase in micronucleus formation, suggesting possible defects in DNA replication or mitotic function (Fig. 4c). We did not observe obvious defects in chromosome alignment in knock-in cells, but overall time in mitosis was reduced in the knock-in clones relative to the WT cells (Fig. 4d). Furthermore, we found that upon nocodazole treatment, both the knock-in cells as well as the inducing cells expressing the AA mutant had a shorter mitotic duration compared to WT and DD-expressing cells (Fig 4e-f). Similar results were obtained using other compounds to arrest cells in mitosis, including the kinesin inhibitor S-trityl-L-cysteine (STLC) and the APC/C inhibitors proTAME and apcin (Extended Data Fig 4c). Since UBE2H interacts with CTLH via its unstructured C-terminal domain, we sought to exclude any potential effects of the C-terminal HA tag. Therefore, we also analyzed cells that express untagged UBE2H. The result showed that cells expressing the AA mutant had a shorter mitotic duration compared to WT and DD-expressing cells, which is consistent with the results using C-terminally HA-tagged UBE2H (Extended Data Fig 4d). Together, these results demonstrate that the non-phosphorylatable UBE2H mutant accelerates mitotic exit and promotes mitotic defects such as micronucleation.

**Fig. 4.**
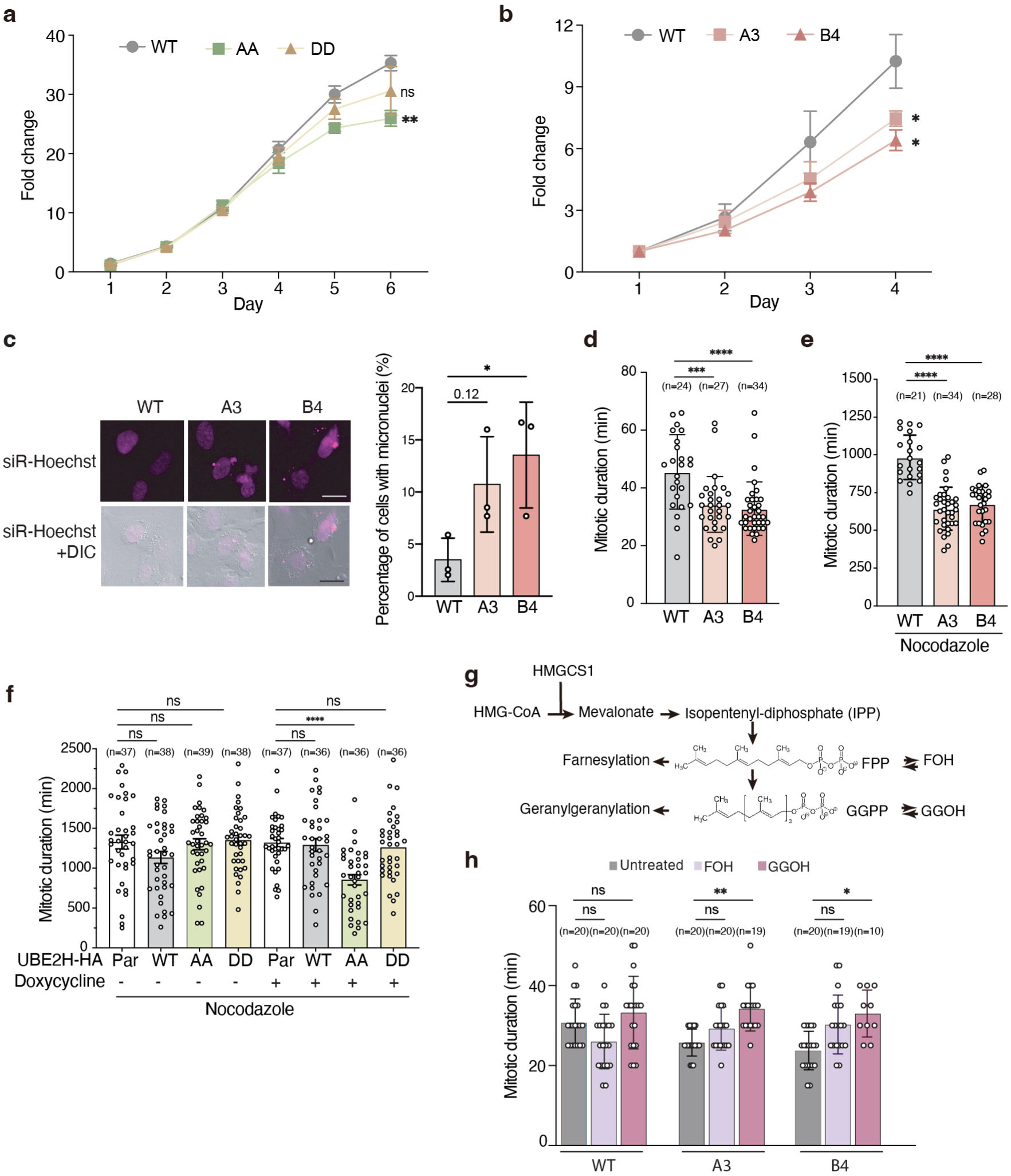
Cells that cannot phosphorylate UBE2H show proliferation and mitotic defects. **a.** Cell viability assays in hTERT-RPE1 cells stably expressing doxycycline-inducible UBE2H constructs (WT, AA and DD) and induced with doxycycline for 6 days. Quantification represents mean ± s.d. from three independent biological replicates. **b.** Cell viability assays were performed using the WT cells and knock-in clones for 4 days. Quantification represents mean± s.d. from three independent biological replicates. **c.** Representative images and quantification of micronucleus formation in WT cells and Clones A3, B4. Scale bar, 10 *µm.* **d.** Mitotic duration of asynchronous WT cells and knock-in clones. Cells were imaged every 3 mins by wide-field time-lapse microscopy for 24 h. **e.** Mitotic arrest duration of asynchronous WT, A3 and B4 cells treated with nocodazole (300 nM) and imaged every 10 mins by wide-field time-lapse microscopy for 48 h. **f.** Mitotic arrest duration of hTERT-RPE1 cells stably expressing doxycycline-inducible UBE2H constructs (WT, AA and DD) were treated with nocodazole (600 nM) and imaged every 10 mins by wide-field time-lapse microscopy for 48 h. **g.** Schematic of the mevalonate pathway highlighting the structures of farnesol (FOH) and geranylgeraniol (GGOH). **h.** Mitotic duration of WT cells and knock-in clones treated with GGOH (50 µM) or FOH (50 µM). For **d-f, h,** Quantification was performed on the number of cells indicated on the plot. Each point represents an individual cell’s mitotic duration, measured as the time from nuclear envelope breakdown (NEB) to division, slippage, or cell death. Error bars indicate mean± s.d. p-values were calculated by one-way ANOVA. **** = p < 0.0001. *** = p < 0.001. ** = p < 0.01. * = p < 0.05. ns = not statistically significant.

Earlier we noted that UBE2H activation can promote the degradation of the CTLH substrate HMGCS1 (Fig. 3a-c). HMGCS1 catalyzes the conversion of acetyl-CoA to HMG-CoA, leading to the production of farnesol (FOH) and geranylgeraniol (GGOH). These molecules are required for protein prenylation (farnesylation and geranylgeranylation) (Fig. 4g). Prior work has shown that expression of kinetochore proteins and mitotic regulators is reduced by inhibition of HMG CoA-reductase and rescued by GGOH^33^. We hypothesized that reduced HMGCS1 levels may lead to defective prenylation, contributing to the mitotic defects observed in UBE2H-AA-expressing cells. We therefore tested whether the downstream products of HMGCS1 could rescue the mitotic defects. Indeed, supplementation with GGOH, a product of the HMGCS1 pathway, significantly rescued the mitotic duration defect in knock-in cells (Fig. 4h). In contrast, FOH showed little effect. These findings suggest that accelerated degradation of HMGCS1 in cells expressing hyperactive UBE2H may contribute to mitotic defects and accelerated mitotic exit. Interestingly, HMGCS1 accumulates in WT cells arrested in mitosis, and this effect was abrogated in the knock-in cells, suggesting that HMGCS1 might be important in mitosis and need to be protected from degradation by inactivating UBE2H (Extended Data Fig. 4e). Taken together, these findings indicate that constitutive activation of UBE2H disrupts normal mitotic progression, perhaps due to inappropriate degradation of HMGCS1. The resulting decrease in protein prenylation may underlie the observed mitotic defects, highlighting the importance of tightly regulated UBE2H activity for accurate cell division.

### Non-phosphorylatable UBE2H expression reduces abundance of CTLH components and substrates

Since the UBE2H-AA mutant is hyperactive in its ability to form ubiquitin thioesters relative to the WT protein, we hypothesized that expression of UBE2H-AA would alter the cellular proteome relative to its WT counterpart. To test this hypothesis, we compared the proteome profiles of parental cells to those expressing doxycycline-inducible UBE2H-WT or the UBE2H-AA mutant. Proteomic analyses were performed on both asynchronous and mitotically-arrested cells (Fig. 5a, Extended Data Fig. 5a, Supplementary Table 1-3). We quantified the fold change of over 8100 proteins in all three cell groups and identified the proteins that were significantly downregulated upon doxycycline induction in either mitosis or interphase (FC >1.25, p < 0.05, Peptides > 1) in each condition. As expected, there were few changes to the proteome in parental cells upon doxycycline addition. In contrast, there were 14 proteins downregulated in asynchronous WT cells and 3 proteins downregulated in mitotic cells. By comparison, a greater number of proteins were significantly downregulated in asynchronous (35 proteins) and mitotic (29 proteins) AA cells (Fig. 5a right, Extended Data Fig. 5b). Notably, there was some overlap between proteins downregulated in WT and AA cells, but there was no overlap in downregulated proteins between either of these groups and the parental cells (Extended Data Fig. 5c). These results indicate that overexpression of AA has a more pronounced impact on the cellular proteome than WT, consistent with our hypothesis that the non-phosphorylatable mutant of UBE2H is more active.

**Fig. 5.**
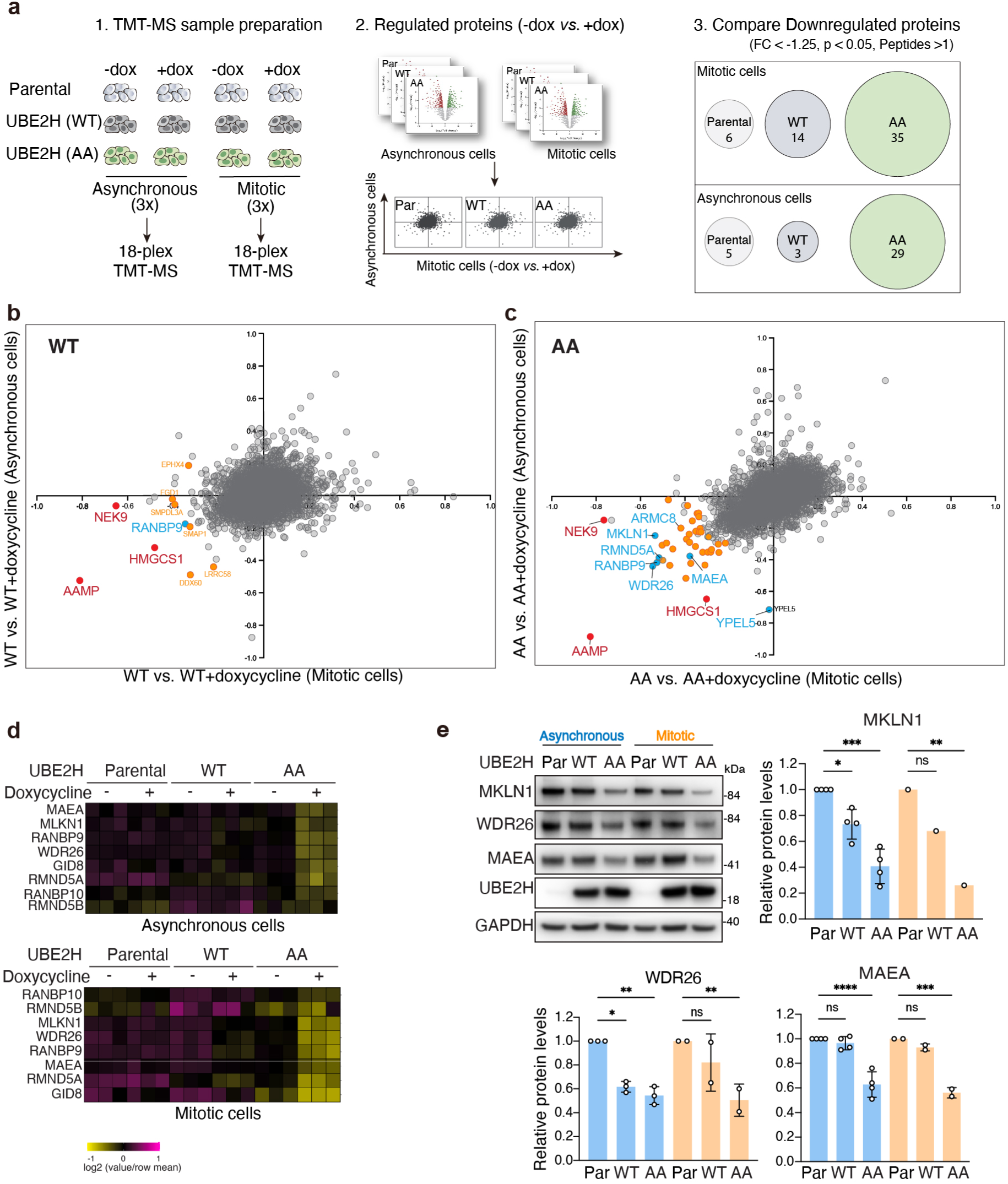
Non-phosphorylatable UBE2H expression reduces abundance of CTLH components and substrates. **a.** Schematic of the TMT-based mass spectrometry workflow to identify proteins regulated by UBE2H (WT and AA) in asynchronous or mitotic cells in three independent biological replicates. (1) hTERT-RPE1 cells stably expressing UBE2H (WT and AA) were cultured in medium with or without doxycycline for 24 h to induce UBE2H expression. Mitotic cells were subsequently collected after an 18 h nocodazole (600 nM) treatment by manually shaking off cells from attached cultures, whereas asynchronous cells were harvested by trypsinization without nocodazole treatment at the same time point. Cell lysates were then subjected to Tandem Mass Tag (TMT)-SPS-MS3 mass spectrometry analysis. (2) Volcano plots of whole cell proteomic analysis comparing effects of doxycycline addition to parental cells or UBE2H (WT, AA)-ex-pressing cells in asynchronous or mitotic conditions. The two-dimensional scatter plots summarize differential changes in asynchronous cells compared with mitotic cells. (3) The specific downregulated proteins in each group were defined as those FC <-1.25, p<0.05 and number of peptides >1. b-c. Two-dimensional plots of TMT-MS analysis comparing the proteins downregulated by UBE2H-WT **(b)** or UBE2H-AA **(c)** in asynchronous and mitotic cells. HMGCS1, AAMP and NEK9 are highlighted in red, and the components of the CTLH complex are highlighted in blue. **d.** Heatmap of the proteomic data curated for components of CTLH E3 ligase. **e.** Western blots showing MKLN1, WDR26, MAEA protein levels in cells overexpressing UBE2H (WT, AA) in asynchronous and mitotic conditions. Quantification represents mean ± s.d. from the indicated independent biological replicates. Statistical significance of differences between groups was determined with one-way ANOVA analysis.****= p < 0.0001,*** = p < 0.001. ** = p < 0.01. * = p < 0.05. ns = not statistically significant.

Next, we compared the degree of downregulation of proteins expressing UBE2H-WT or UBE2H-AA in asynchronous or mitotic conditions (Fig. 5b-c, Extended Data Fig. 5d), to determine whether downregulation depended on cell cycle phase. In both WT cells and AA cells, we found that the known CTLH substrate, HMGCS1, was downregulated in both asynchronous and mitotic cells. Furthermore, Angio-associated Migratory Protein (AAMP) was markedly downregulated in both WT and AA cells under asynchronous and mitotic conditions. In contrast, we found that the serine/threonine kinase NEK9 was downregulated in mitotic cells but not asynchronous cells. Finally, we observed that components of the CTLH complex including RANBP9, RMND5A, MAEA, ARMC8, MKLN1, WDR26 and YPEL5 (Fig. 5d, Supplementary Table 4) were downregulated, to a greater extent in the AA cells compared to the WT cells. Interestingly, YPEL5 was downregulated in asynchronous cells but not mitotic cells. We verified these results by western blotting (Fig. 5e). These results support the hypothesis that UBE2H-AA is more active, especially in its potential to downregulate activity of the CTLH complex.

In addition, we compared the proteomic profiles of knock-in cells (clones A3 and B4) to their parental counterparts in asynchronous and mitotic populations (Extended Data Fig. 5e-f, Supplementary Table 1-3). Protein levels were highly concordant between the two clones in both asynchronous (r = 0.9277, p < 0.0001) and mitotic cells (r = 0.9123, p < 0.0001). Like the inducing cells, several components of the CTLH complex (MKLN1, MAEA) were downregulated in the knock-in cells. Moreover, HMGCS1, NEK9 and AAMP also appeared among the downregulated proteins (Extended Data Fig. g-h). Although many more proteins were altered in the knock-in cells than the inducing cells, we speculate that broader changes may arise during the selection of clones in the context of the growth disadvantage caused by expression of hyperactive UBE2H. Taken together, these proteomic analyses are consistent with the idea that the non-phosphorylatable mutant of UBE2H exhibits higher ubiquitin-conjugating activity than the wild-type enzyme, which in turn enhances both CTLH activity and promotes its downregulation.

### CTLH targets NEK9 for degradation in mitosis

We identified NEK9 and AAMP as two strongly-downregulated proteins in cells expressing non-phosphorylatable UBE2H. We next sought to determine whether this downregulation was a direct result of CTLH-dependent degradation. Interestingly, NEK9 appeared to be downregulated only during mitosis in cells overexpressing either UBE2H-WT or UBE2H-AA, with the latter showing a stronger effect (Fig. 6a). We confirmed these results by western blotting (Fig. 6b). Additionally, when we used other mitotic inhibitors to arrest cells in mitosis, NEK9 was similarly downregulated by UBE2H overexpression (Extended Data Fig. 6a). We performed a time-course analysis and found that downregulation is detectable beginning about 4 hours after mitotic entry (Extended Data Fig. 6b). Furthermore, we verified the protein levels of NEK9 in knock-in cells and found that NEK9 was also downregulated in the two AA knock-in clones during mitosis but not in interphase (Fig. 6c-d).

**Fig. 6.**
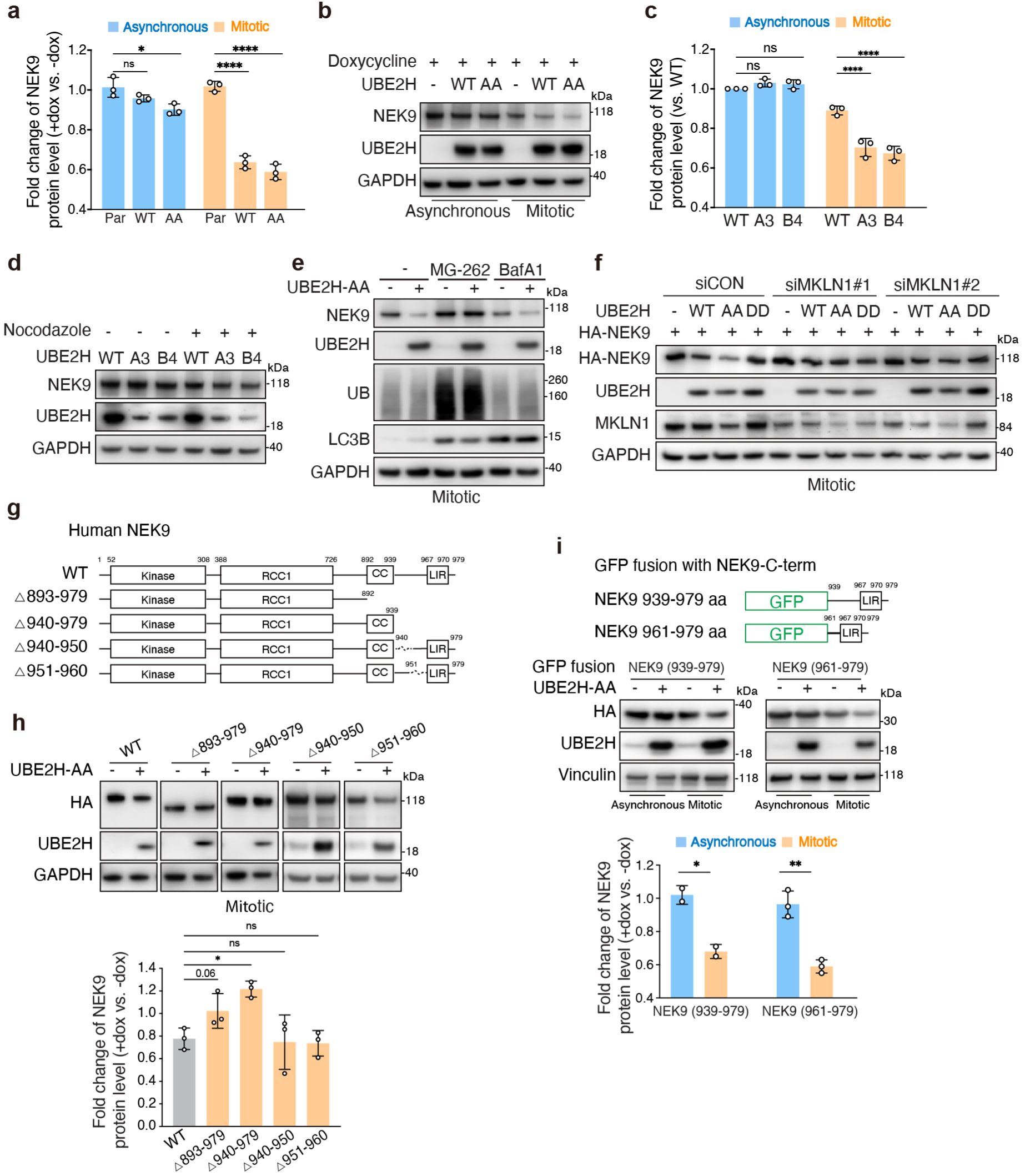
CTLH targets NEK9 for degradation in mitosis. **a.** Downregulation of NEK9 measured by TMT-MS in parental and UBE2H (WT, AA) hTERT-RPE1 cells. Quantification represents mean ± s.d. from three independent biological replicates. **b.** Western blots showing NEK9 protein levels in hTERT-RPE1 cells expressing UBE2H-AA in asynchronous and mitotic conditions. **c.** Downregulation of NEK9 measured by TMT-MS in WT cells and knock-in clones (A3 and B4). Quantification represents mean± s.d. from three independent biological replicates. **d.** Western blots showing NEK9 protein levels in WT cells and knock-in clones in asynchronous and mitotic conditions. **e.** Western blots showing NEK9 protein levels in mitotic UBE2H-AA-expressing hTERT-RPE1 cells after treatment with MG262 (10 µM) or bafilomycin A1 (BafA1, 1 µM) in the presence or absence of doxycycline. **f.** Western blots showing exogenous NEK9 protein levels in mitotic parental and UBE2H (WT, AA, DD) hTERT-RPE1 cells transfected with control siRNA (siCON) or two independent MKLN1 siRNAs. **g.** Schematic representation of human NEK9 constructs (WT, Δ893-979, Δ940-979, Δ940-950, Δ951-960). **h.** Western blots showing NEK9 (WT, Δ893-979, Δ940-979, Δ940-950, Δ951-960) protein levels in mitotic UBE2H-AA-expressing hTERT-RPE1 cells in the presence or absence of doxycycline. Quantification represents mean± s.d. from three independent biological replicates. i. Schematic of GFP fusion with NEK9-C-terminal fragments (top). Western blots showing GFP-NEK9 C-terminal fusion protein levels upon UBE2H (AA) induction in mitotic conditions (middle). Quantification represents mean ± s.d. from three independent biological replicates (bottom). Statistical significance of differences between groups was determined with one-way ANOVA analysis (a, c and h) or two-tailed unpaired Student’s t-test (i). **** = p < 0.0001. ** = p < 0.01. * = p < 0.05. ns = not statistically significant.

To investigate the mechanism underlying NEK9 downregulation by UBE2H, we first examined whether it was due to proteasomal degradation. We found that the proteasome inhibitor MG262 rescued the downregulation of both endogenous and exogenous NEK9 induced by UBE2H-AA (Fig. 6e, Extended Data Fig. 6c). In contrast, the lysosome inhibitor bafilomycin A1 (BafA1) showed no effect. Next, to determine whether the CTLH complex mediates the downregulation of NEK9, we used chemical inhibitors or siRNA to transiently inhibit the components of CTLH complex in UBE2H-overexpressing cells. First, we used PFI-7, a compound that binds to the substrate receptor GID4 and competes with substrates for binding^34^. We found that PFI-7 had no effect, indicating that NEK9 is not recognized by the GID4 substrate receptor (Extended Data Fig. 6d). Next, we knocked down MKLN1 or WDR26 and found that knockdown of MKLN1 blocked the degradation of both endogenous and exogenous NEK9 (Fig. 6f, Extended Data Fig. 6e). In contrast, knockdown of WDR26 had no effect (Extended Data Fig. 6f-g). Thus, our findings indicate that NEK9 is downregulated by CTLH-mediated proteasomal degradation that depends on the CTLH complex component MKLN1, but not on GID4 or WDR26.

Next, we sought to identify the degron required for NEK9 degradation in mitosis. NEK9 consists of an N-terminal kinase domain, followed by autoinhibitory regulator of chromosome condensation 1 (RCC1)-repeats, a coiled-coil domain and a conserved LIR motif within a long C-terminal disordered region. We generated a series of C-terminal truncation mutants (△893-979, △940-979, △940-950, △951-960) and found that two of them (△893-979, △940-979) were resistant to CTLH/UBE2H-mediated degradation, suggesting that 19 residues in the C-terminal region (961-979) are necessary for NEK9 degradation (Fig. 6g-h, Extended Data Fig. 6h). To test sufficiency, we fused the NEK9 C-terminal fragments (aa 939-979 or 961-979) to GFP and found that both fusion proteins were downregulated by UBE2H-AA expression in mitosis, demonstrating that the C-terminus of NEK9 (aa 961–979) is sufficient to confer mitosis-specific degradation to a heterologous protein (Fig. 6i). In summary, these findings demonstrate that overexpression of hyperactive UBE2H can induce NEK9 degradation in mitosis, and that this process requires the CTLH complex component MKLN1 and depends on a degron located within C-terminus of NEK9.

### CTLH-MKLN1 targets AAMP for degradation via DR-like C degron

Next, we sought to determine whether the downregulation of AAMP was a direct result of CTLH-dependent degradation. Among the proteins that were downregulated preferentially in the presence of non-phosphorylatable UBE2H, AAMP showed the most dramatic change in abundance (Fig. 7a). We validated by western blotting that AAMP can be downregulated by UBE2H-WT and AA in both asynchronous and mitotic cells (Fig. 7b). In addition, we verified that the level of AAMP was also downregulated in the two knock-in clones in both asynchronous and mitotic populations (Fig. 7c).

**Fig. 7.**
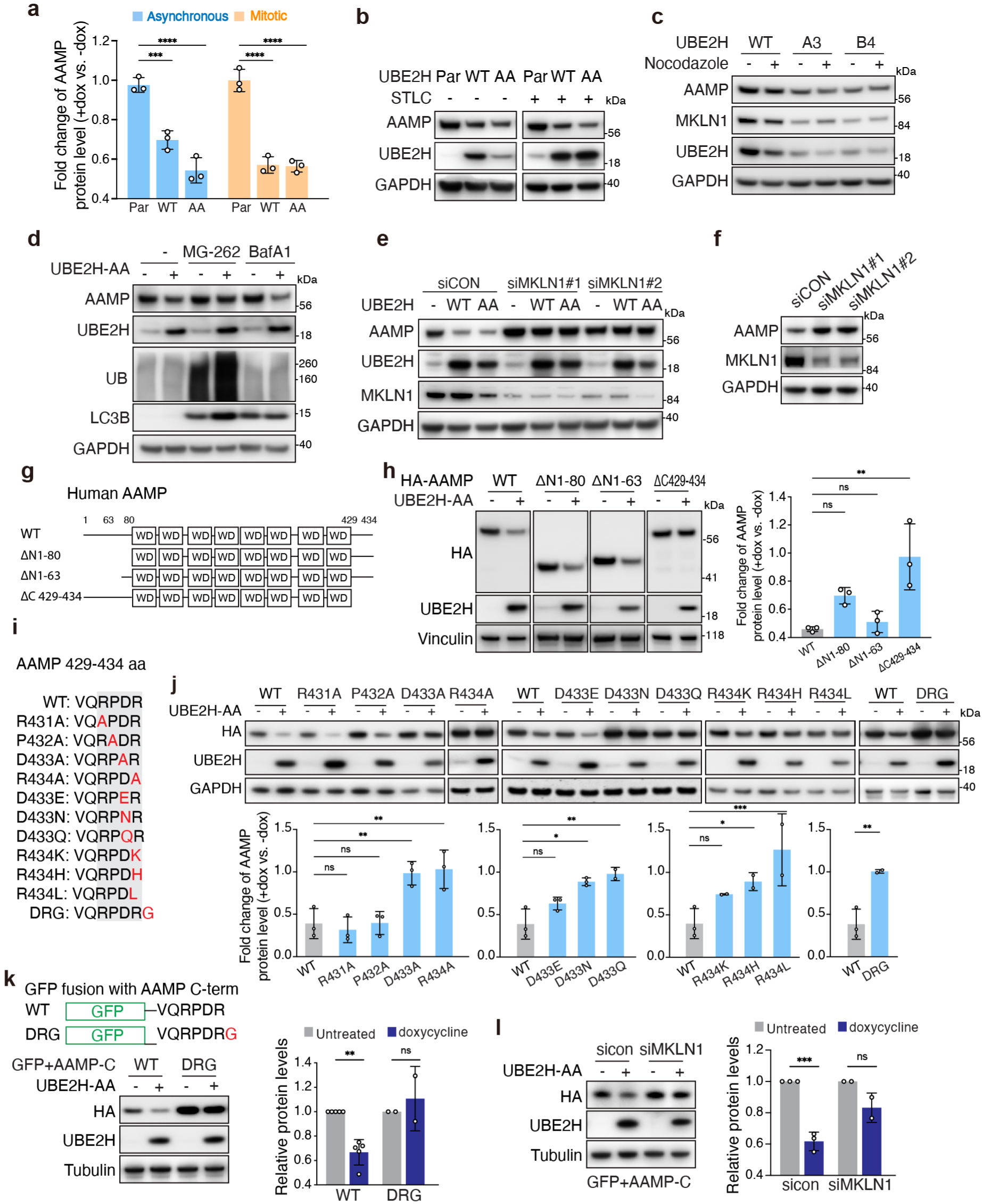
CTLH-MKLN1 targets AAMP for degradation via DR-like C degron. **a.** Downregulation of AAMP as measured by TMT-MS in parental and UBE2H (WT, AA) RPE1 cells. Quantification represents mean ± s.d. from three independent biological replicates. **b.** Western blots showing AAMP downregulation in hTERT-RPE1 cells upon induction of UBE2H (WT, AA) in asynchronous and mitotic conditions. **c.** Western blots showing AAMP downregulation in A3 and B4 hTERT-RPE1 cells in asynchronous and mitotic conditions. **d.** Western blots showing AAMP protein levels in UBE2H-AA-expressing hTERT-RPE1 cells after treatment with MG-262 (10 µM) or BafA1 (1 µM) in the presence or absence of doxycycline. e. Western blots showing AAMP protein levels in parental and UBE2H (WT, AA) hTERT-RPE1 cells transfected with control or MKLN1 siRNAs. **f.** Western blots showing AAMP protein levels in hTERT-RPE1 cells transfected with control or MKLN1 siRNAs. **g.** Sequence schematic of human AAMP and derivatives. **h.** Western blots showing levels of AAMP and derivatives in UBE2H-AA-expressing hTERT-RPE1 cells in the presence or absence of doxycycline. Quantification represents mean± s.d. from three independent biological replicates. i. AAMP C-terminal amino acid sequence (aa 429-434). The conserved regions are highlighted in gray. The mutations are colored in red. **j.** Western blots showing AAMP (WT, mutants) protein levels in UBE2H-AA-expressing hTERT-RPE1 cells in the presence or absence of doxycycline. The fold change of AAMP (WT, mutants) protein levels is quantified (+dox vs -dox). Quantification represents mean± s.d. from three independent biological replicates. k. Diagram of GFP fusions with AAMP-C-termini and western blots showing protein levels upon UBE2H-AA induction. Quantification represents mean± s.d. from three independent biological replicates. I. Western blots showing protein levels of GFP fusions in UBE2H-AA-expressing hTERT-RPE1 cells transfected with control siRNA or siRNA targeting MKLN1 (siMKLN1#1). Quantification represents mean ± s.d. from three independent biological replicates. For quantification results, p-values were calculated by one-way ANOVA. **** = p < 0.0001. *** = p < 0.001. ** = p < 0.01. * = p < 0.05. ns = not statistically significant.

To determine whether AAMP downregulation by UBE2H was due to proteasomal degradation, we treated cells with proteasome inhibitor MG262 or lysosome inhibitor BafA1. The results showed that MG262, but not than BafA1, rescued the downregulation of both endogenous and exogenous AAMP induced by UBE2H-AA, suggesting that UBE2H-AA mediated AAMP degradation in a proteasome-dependent manner (Fig. 7d, Extended Data Fig 7a). Next, to determine whether the CTLH complex mediates the downregulation of AAMP, we examined the effects of knock down of MKLN1 and found that it blocked the degradation of AAMP in the presence of UBE2H-AA (Fig. 7e). Downregulation of an alternative CTLH adapter protein, WDR26, had no effect (Extended Data Fig. 7b). Furthermore, knockdown of MKLN1, but not WDR26, resulted in the accumulation of AAMP even in the absence of overexpression of UBE2H-AA, suggesting that MKLN1 is a critical regulator of AAMP under normal conditions (Fig. 7f, Extended Data Fig. 7c). In contrast, the GID4 inhibitor PFI-7 had no effect, suggesting AAMP was recognized independently of GID4 (Extended Data Fig. 7d). Thus, these results indicate that the CTLH complex regulates AAMP stability in a manner that requires the adaptor MKLN1, but not GID4 or WDR26.

We next performed a truncation analysis of AAMP to identify potential degrons. AAMP is a 434-amino acid protein that is predicted to contain a structured beta-propeller domain consisting of WD40 repeats, which is flanked by largely unstructured N- and C-terminal regions. We started by deleting the unstructured regions, constructing three truncation mutants of AAMP (△N1-63, △N1-80 and △C429-434 (Fig. 7g). Only △C429-434 showed resistance to CTLH/UBE2H-mediated degradation, suggesting the C-terminus of AAMP (429-434aa) was necessary for its degradation (Fig. 7h). This region consists of six amino acids “VQRPDR”, in which the “PDR” motif is highly conserved among vertebrates (Extended Data Fig. 7e).

Mutational analysis of the “RPDR” motif revealed that substitutions within the C-terminal “DR” residues abolished CTLH/UBE2H-mediated degradation. Substitutions at D433 indicated a requirement for a negative charge at this position: D433E remained degradable, whereas D433N and D433Q stabilized AAMP. Mutations at R434 suggested a requirement for a basic residue with an appropriate side chain: R434K was still degraded, while R434H and R434L blocked degradation. Finally, inserting an additional glycine at the C-terminus abolished degradation, indicating that the DR sequence must be positioned at the C-terminus for degron function (Fig. 7i-j). In addition, GFP fused to AAMP (429-434aa) was also degraded upon expression of UBE2H-AA and this degradation was rescued either by adding an extra glycine residue at the C-terminus or by knock down of MKLN1, demonstrating that the C-terminus of AAMP is sufficient to confer degradation on a heterologous protein (Fig. 7k). Furthermore, knock down of MKLN1 blocked the degradation of this fusion protein in the presence of UBE2H-AA, suggesting that it is regulated similarly to the full-length protein (Fig. 7l). Taken together, these results suggest a novel C-degron recognition mechanism in which the “DR” tail mediates degradation of AAMP by UBE2H/CTLH-MKLN1.

### MKLN1 recognizes a DR-like C-degron as a CTLH substrate receptor

To identify the substrate receptor for AAMP, we performed a proteomic analysis of AAMP-associated proteins in cells overexpressing AAMP or two stable mutants (△C and D433A) to identify the AAMP-interacting partners that depended on its “DR” tail (Fig. 8a). By comparing peptide spectral counts, we identified 53 proteins that specifically interacted with AAMP-WT, including four CTLH subunits (MKLN1, MAEA, RMND5A and ARMC8). Among these, MKLN1 ranked first based on the number of peptide spectral counts (Fig. 8b, Supplementary Table 5). In addition, we applied AlphaFold 3 (AF3) to predict the interaction between AAMP and all CTLH components^35^. MKLN1 showed the strongest predicted interaction with AAMP, with an ipTM of 0.54 and a pTM of 0.61 (Fig. 8c, Supplementary Table 6). In contrast, AAMP mutants with substitutions in the C-terminal residues showed much lower predicted interaction scores (ipTM 0.20, pTM 0.48) (Extended Data Fig. 8a). These results suggested that MKLN1 might function as the substrate receptor for AAMP, which requires the C-terminal “DR” tail of AAMP for recognition. In analyzing the AF3 model, AAMP was predicted to interact with the Kelch-repeat domain of MKLN1 via its C-terminus (Fig. 8d). Notably, D433 and R434 were positioned adjacent to oppositely charged surface residues on MKLN1, suggesting possible electrostatic interactions (Fig. 8e). Meanwhile, we noticed that another reported substrate of CTLH, Zinc finger MYND domain-containing protein 19 (ZMYND19)^24^, has a C-terminal “ER” tail resembling that of AAMP. AF3 prediction indicated that ZMYND19 similarly interacted with MKLN1 Kelch-repeat domain through its C-terminus, with upstream negatively charged residues E222 and E224 likely enhancing the interaction (Extended Data Fig. 8b). In addition, HMGCS1 also has a similarly charged C-terminal “EH” tail. Although the upstream “VI” motif was reported to be required for HMGCS1 degradation^25^, HMGCS1 also interacted with the Kelch-repeat Domain of MKLN1 via its C-terminus. These observations suggest that MKLN1 recognition of “DR”-like C-terminal tail may represent a general recognition mechanism for CTLH substrates.

**Fig. 8.**
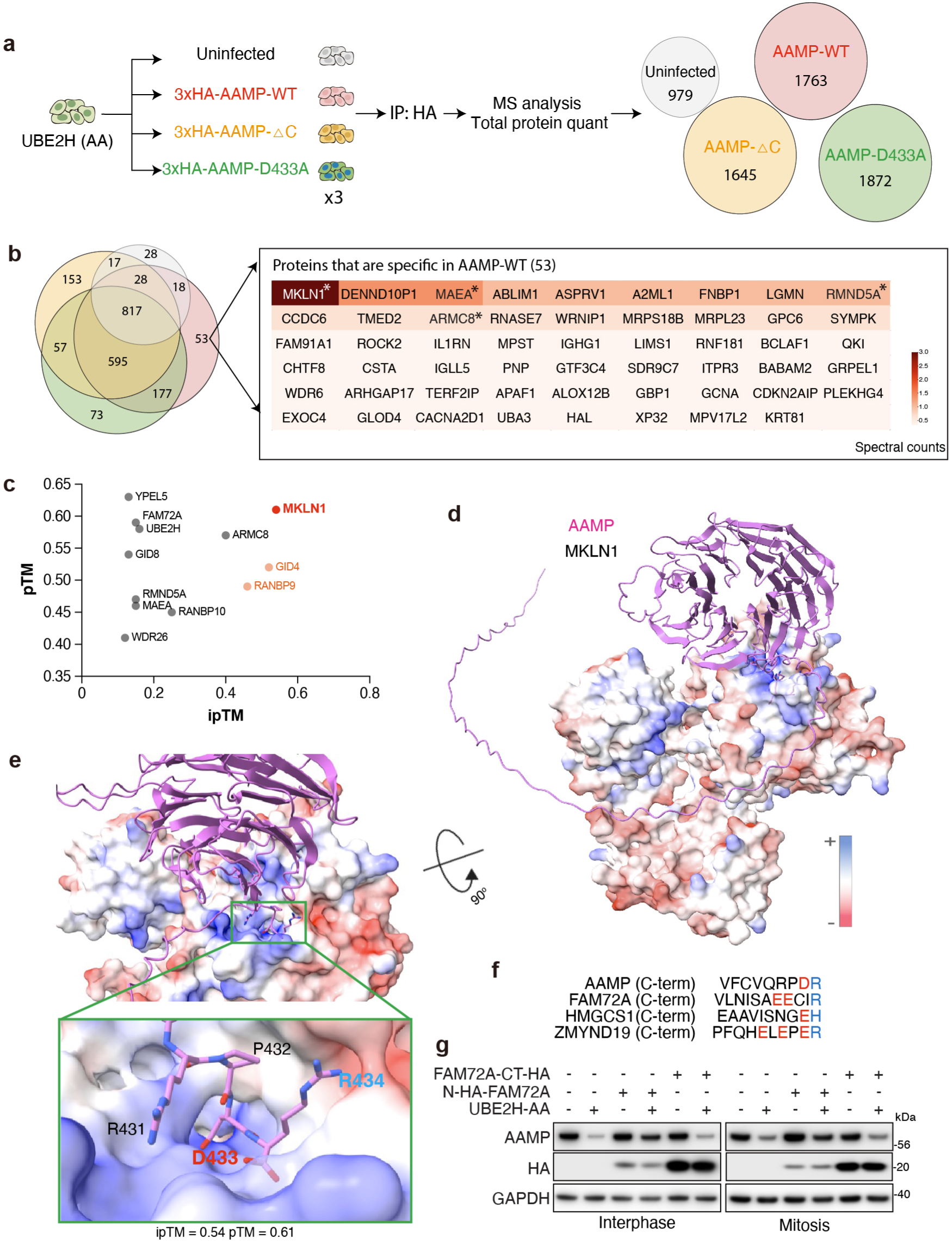
MKLN1 recognizes DR-like C-degron as a CTLH substrate receptor. **a.** Schematic of the mass spectrometry workflow to identify the AAMP-associated proteins in cells overexpressing wild-type AAMP or two stable AAMP mutants (ΔC429-434 and D433A) in three biological replicates. **b.** Venn diagram showing overlap between proteins and those that specifically interacted with AAMP-WT. Proteins are ordered according to their spectral counts. **c.** Predicted interactions between AAMP and all CTLH components using AlphaFold 3 (AF3), showing ipTM (x-axis) versus pTM (y-axis) scores. **d-e.** AF3 models for AAMP (pink)-MLKN1 interaction, with MKLN1 colored by electrostatic potential surface. Residues at the extreme AAMP C-terminus interact with a groove formed by the blades of MKLN1 β-propeller. ipTM scores for pairwise combinations are indicated. **f.** AAMP C-terminal amino acid sequence (aa 429-434) showing a putative degron (acidic residues in red, basic residues in blue), aligned with FAM72A, HMGCS1and ZMYND19 C-terminal sequence, revealing a consensus motif with a terminal basic residue preceded by acidic residues. **g.** Western blots showing AAMP protein levels in UBE2H-AA-expressing hTERT-RPE1 cells upon expression of N-terminally or C-terminally tagged FAM72A.

We further examined FAM72A, an adaptor that interacts with MKLN1 to target the substrate UNG2 for degradation^36^. FAM72A carries a C-terminal “IR” tail that similarly interacted with the MKLN1 kelch repeat domain through its C-terminus, with upstream acidic residues E145, E146 likely strengthening the interaction (Fig. 8f, Extended Data Fig. 8c). Because FAM72A and AAMP appeared to bind the same MKLN1 domain, we hypothesized that FAM72A might compete for degradation with substrates like AAMP, ZMYND19, and HMGCS1. To test this, we constructed both C-terminally and N-terminally HA/FLAG-tagged FAM72A constructs and introduced them into inducible UBE2H-AA cells. As expected, endogenous AAMP was degraded in the presence of UBE2H-AA. Interestingly, N-terminally tagged FAM72A stabilized AAMP, whereas C-terminally tagged FAM72A did not (Fig. 8g), as expected if the C-terminus of FAM72A is required for binding MKLN1. These results support the hypothesis that FAM72A competes with AAMP for binding to MKLN1 and are consistent with a model in which the C-terminus of FAM72A interacts with MKLN1 to compete for binding of other substrates such as AAMP. Taken together, these results indicate that MKLN1 functions as a substrate receptor that recognizes the substrates containing “DR”-like C-terminal tail, whereas FAM72A acts both as an adaptor of MKLN1 for its substrate UNG2, and as a competitor for substrates like AAMP, ZMYND19 and HMGCS1.

## Discussion

The mammalian CTLH complex appears responsive to cell-cycle and nutrient cues, yet the specific mechanisms remain incompletely understood. Our findings identify phosphorylation of the cognate E2 enzyme, UBE2H, as such a control point: CDK and mTOR phosphorylate UBE2H at two conserved N-terminal serine residues (S3/S5) and reduce UBE2H charging with ubiquitin, thereby limiting the pool of active E2 available to the CTLH ligase. mTOR and CDKs both recognize proline-directed serine/threonine motifs (S/T-P), with CDK-dependent phosphorylation augmented in the context of an extended S/TPXK/R consensus^37^. While S3 conforms to the minimal S/T-P motif, S5 matches the extended CDK consensus. Cells expressing non-phosphorylatable UBE2H display enhanced UBE2H charging during interphase and mitosis, promoting CTLH-mediated substrate degradation and establishing UBE2H phosphorylation as a switch that tunes CTLH activity across different contexts. By taking advantage of hyperactive UBE2H, we identify two novel substrates of the CTLH complex and define a distinct C-degron recognition mode, in which a DR-like C terminal tail is recognized by MKLN1, supporting a direct substrate receptor function for MKLN1 within the CTLH complex. Together, these findings identify a CDK/mTOR–UBE2H axis that gates CTLH-dependent degradation by controlling E2-ubiquitin thioester availability and establish E2 phosphorylation as a proximal control point for context-dependent ubiquitylation (Extended Data Fig. 8d).

Mechanistically, N-terminal phosphorylation of the E2 enzyme UBE2H inhibits CTLH E3 ligase activity by directly suppressing its ability to be charged by E1, thereby reducing formation of the UBE2H∼Ub thioester, limiting the amount of active E2 that can be engaged by the CTLH ligase. This result is consistent with the prior observation that the presence of an acidic residue or phosphorylation in the E2 N-terminal region may inhibit its ability to be charged by E1^9^. Notably, among known E2s, UBE2H thus appears to be susceptible to a high degree of phospho-regulation: CK2-mediated phosphorylation of UBE2H in its C-terminal region has been shown promote its association with CTLH and its ability to ubiquitylate substrates^31^. C-terminal phosphorylation may establish a permissive baseline for UBE2H-CTLH engagement, whereas CDK/mTOR-mediated N-terminal phosphorylation provides a dynamic brake that couples cell cycle and nutrient cues to the availability of charged E2. At the same time, inhibitory phosphorylation of the N-terminus of UBE2H may be important for preserving CTLH activity once substrates are cleared. In this scenario, a mechanism to inactivate UBE2H while it is associated with CTLH may be required to preserve E3 activity, which is consistent with our observation that CTLH component expression declines more in UBE2H-AA-expressing cells compared to UBE2H-WT expressing cells.

Our results indicate that mTOR and CDK both have the capacity to inactivate a common target, UBE2H. A similar functional relationship between mTOR and CDK1 has been previously identified as a key regulatory strategy for autophagy, where mTOR restrains autophagy during interphase, and CDK1 restrains autophagy in mitosis. Because CDK1 activation leads to mTOR inhibition by phosphorylation of Raptor, CDK1 is required to take over for mTOR to maintain inhibition of autophagy during mitosis^37,38^. Our results suggest that mTOR and CDK may operate in a comparable “handoff” mode for UBE2H across the cell cycle. In addition, our findings offer a mechanistic explanation for the previously observed activation of CTLH-dependent degradation by mTOR inhibitors in asynchronous cell populations^25–27^. Our results show that cells expressing non-phosphorylatable UBE2H phenocopy mTOR inhibition and are largely insensitive to further mTOR inhibition with respect to CTLH-dependent degradation, placing UBE2H phosphorylation downstream of mTOR. Interestingly, it was recently shown that the CTLH substrate ZYMND19 and receptor MKLN1 can block mTORC1 activation at the lysosome when CTLH is inactive^24^. Therefore, our findings suggest that mTOR-mediated inhibition of UBE2H may complete a negative feedback loop to enable homeostasis of mTOR activity during interphase.

Our results show that proper mitotic progression appears to require inhibition of UBE2H-CTLH function. Cells expressing non-phosphorylatable UBE2H exhibit impaired proliferation and mitotic fidelity, including accelerated mitotic exit and micronucleus formation. Interestingly, UBE2H protein levels decrease during mitosis (Extended Data Fig. 4d), which may synergistically work with phosphorylation to further limit the available UBE2H∼Ub pool and prevent aberrant CTLH activity during cell division. Two CTLH substrates of interest in this study provide a plausible mechanistic entry point for these phenotypes. First, supplementation with HMGCS1 downstream metabolites rescued mitotic defects in UBE2H-AA-expressing cells, indicating that reduced flux in the mevalonate pathway may limit geranylgeranylation required for proper mitosis. Second, inhibition of UBE2H in mitosis may help stabilize NEK9, a recognized regulator of spindle dynamics and centrosome segregation^39,40^. We have shown that a region in the C-terminus of NEK9 is necessary for its degradation, but why NEK9 is only degraded in mitosis remains unclear. Together, these observations support a model in which UBE2H is negatively regulated to maintain key metabolic pathways and NEK9-dependent function for proliferation and accurate chromosome segregation.

By leveraging hyperactive UBE2H, we uncovered AAMP as a CTLH substrate with MKLN1 functioning as the substrate receptor. Notably, AAMP was previously identified as a cytoplasmic interactor of CTLH ^41^. In addition, prior work suggested that MKLN1 might function as a substrate receptor; however, how it contributes to substrate specificity and ubiquitylation remained unclear^18,19,25^. We have discovered a DR-like C-terminal degron class, which expands the CTLH substrate landscape and provides a mechanism for MKLN1-dependent targeting. Degron mapping and GFP fusion assays showed that a C-terminal DR-like tail is both necessary and sufficient for CTLH-dependent degradation. IP–MS proteomics together with AlphaFold3-based modelling suggests that MKLN1 recognizes AAMP by a DR-like C-degron. This model is supported by our findings that FAM72A can compete with this interaction. While preparing this manuscript, a preprint reported AAMP and ZMYND19 as CTLH-regulated substrates and proposed MKLN1 involvement in this C-degron pathway^28^. Our data align with this model, and extend it by (i) defining the minimal C-terminal sequence requirements for AAMP by showing that the C-terminal degron is sufficient to confer CTLH-dependent degradation on a heterologous protein, (ii) generalizing the rule into a broader DR-like degron class (including related ER- and DK-like C-termini), and (iii) providing evidence for competitive gating of MKLN1-dependent recognition by FAM72A.

UBE2H appears to act as a dedicated E2 for CTLH, which may explain why regulation at the E2 level is both effective and specific. Consistent with this notion, our proteomic analyses indicate that overexpression of UBE2H does not trigger widespread protein degradation, arguing against broad engagement of other E3 ligases when UBE2H is overexpressed. Since many E2s collaborate with multiple E3s, tuning E3 activity via regulation of E2 charging may not be expected to provide E3-specific control in most cases. This may explain why most E2s do not seem to be phospho-regulated. However, an exception maybe the known stably-associated E2/E3 pair, RAD6A/B (UBE2A/UBE2B)–RAD18, a canonical E2–E3 pair involved in post-replication DNA damage tolerance^42,43^. Interestingly, our screen in *Xenopus* egg extract revealed that UBE2A and UBE2B charging is also reduced in mitosis and both also have N-terminal serine/threonine residues that may be candidate phosphoregulatory sites. It will therefore be interesting to test whether these enzymes are also phospho-regulated in cells to control the rate of RAD18-dependent ubiquitylation. Overall, our findings support a model in which phospho-regulation of E2 charging may be a key mechanism to tune the output of tightly coupled E2–E3 pairs, thereby achieving E3-specific control.

## Limitations

We have identified phosphorylation of UBE2H as an important mechanism for gating CTLH-dependent degradation, but several limitations remain. First, our conclusions are largely based on cultured-cell perturbations (kinase inhibition and mutation of UBE2H). Second, how different cyclin–CDK complexes contribute to UBE2H inactivation during interphase, and the cooperation with mTOR-dependent regulation during G2, S and G2 phases, remains to be explored. Next, although our data support MKLN1-dependent recognition of a DR-like C-terminal degron class and reveal competition by FAM72A, more direct biochemical evidence including binding and *in vitro* reconstructed ubiquitylation assays as well as structural studies will more fully illuminate how CTLH-MKLN1 recognizes this degron. Finally, while we identify new CTLH substrates and a DR-like C-degron rule, the breadth of this degron class, its prevalence within the proteome, and the extent to which additional CTLH receptors (beyond MKLN1 and GID4) contribute to context-specific targeting remain open questions.

## Methods

### Cell lines

HEK293T and HeLa cells were grown in Dulbecco’s modified Eagle’s medium (DMEM). hTERT-RPE1 cells and hTERT-RPE1 cells stably expressing H2B-RFP (a gift from A. Holland) were grown in DMEM/Hams F-12 50/50 mix. All cell lines were supplemented with 10% fetal bovine serum (Gemini Bioproducts LLC) and maintained in a humidified 5% CO2 incubator at 37°C. All cell lines were found to be free of mycoplasma using Mycoplasma Plus PCR assay kit (SouthernBiotech).

### Antibodies

The following antibodies were used: UBE2H (Invitrogen, PA5112141), NEK9 (Proteintech, 11192-1-AP), AAMP (Invitrogen, JE64-70), HMGCS1 (Proteintech, 17643-1-AP), Phospho-S6 Ribosomal Protein (Ser240/244) Antibody (Cell Signaling Technology, 2215S), S6 Ribosomal Protein (5G10) Rabbit Monoclonal Antibody (Cell Signaling Technology, 2217S), MKLN1 (Santa Cruz Biotechnology, sc-398956), WDR26 (Thermo Fisher Scientific, PA555892), MAEA (R&D, AF7288), Cyclin B1 (Santa Cruz Biotechnology, sc-245), CDC27 (BD Biosciences, 610455), Vinculin (Santa Cruz Biotechnology, sc73615), Tubulin (Santa Cruz Biotechnology, sc-32293), GAPDH (Proteintech, 600004-1), HA-Peroxidase (Roche, 12013819001), TSC2 (ABclonal, A19540), IRDye 800CW Goat anti-Rabbit IgG H+L (LI-COR, 926-32211), IRDye 800CW Goat anti-Mouse IgG H+L (LI-COR, 926-32210), Goat anti-Rabbit IgG (H+L) Secondary Antibody, HRP (Invitrogen, 31460), Goat anti-Mouse IgG (H+L) Secondary Antibody, HRP (Invitrogen, 31430).

### Chemicals

The following chemicals were used: doxycycline hyclate (Sigma Aldrich, D9891), RO3306 (AdipoGen Life Sciences, AGCR13515M), (R)-roscovitine (AdipoGen Life Sciences, AG-CR1-0006-M005), nocodazole (Selleckchem, S2775), (+)-S-trityl-L-cysteine (STLC) (Alfa Aesar, L14384), proTAME (Boston Biochem, I-440), apcin (Enamine, T0506-3874), taxol (Sigma-Aldrich, 33069-62-4), thymidine (Sigma-Aldrich, T9250), palbociclib (LC Laboratories, P-7722), torin1 (Cell Signaling, 14379S), rapamycin (RPI, R64500-0.001), MG-132 (Calbiochem, 474790), MG-262 (Apexbio, A8179-1), cycloheximide (Sigma, C7698), geranylgeranyl alcohol (Cayman Chemical, 13272), farnesyl alcohol (Cayman Chemical, 13268), puromycin (Thermo Scientific, 22742-0100), bafilomycin A1 (RPI, B40500-0.001), PFI-7 (Gift from C. H. Arrowsmith, Structural Genomics Consortium), SiR-DNA (SiR-Hoechst*) (Spirochrome, SC007), Pierce protease inhibitor tablet (Thermo Scientific, A32953), phosphatase inhibitor tablet (Thermo Scientific, A32957).

### Plasmids

To generate doxycycline-inducible UBE2H constructs, the hORFEOME UBE2H ORF (Harper laboratory, Harvard Medical School) was cloned into pINDUCER20 (Addgene plasmid #44012, gift from S. Elledge) using the Gateway LR Clonase II system (Invitrogen). An untagged UBE2H version was generated by PCR-based site-directed mutagenesis. For FLAG-HA-tagged constructs, hORFeome entry clones encoding NEK9, AAMP, FAM72A were cloned into the pHAGE-FLAG-HA-NTAP vector (gift from J. W. Harper) using the Gateway LR Clonase II system (Invitrogen). C-terminally HA/FLAG-tagged FAM72A construct was cloned into the pHAGE-FLAG-HA-CTAP vector (gift from J. W. Harper) using the Gateway LR Clonase II system (Invitrogen). Point mutants of UBE2H, NEK9, AAMP were generated by PCR-based site-directed mutagenesis on the corresponding wild-type template. All engineered sequences were confirmed by Sanger sequencing at the entry-clone stage and again after assembly of the final expression vectors. All plasmids and mutagenesis details are listed in Supplementary Table 7.

### Lentivirus Construction

Lentiviral particles were generated in HEK293T cells by co-transfecting the packaging plasmid pPAX2, the envelope plasmid pMD2 (gift from J. W. Harper), and the indicated transfer vector at a 4:2:1 DNA ratio using PEI MAX® Transfection Reagent (Kyfora Bio, 24765) according to manufacturer’s instructions. At 16-18 h post-transfection, HEK293T cells were switched to fresh media supplemented with 1 mM sodium pyruvate. At 48 h post-transfection, lentivirus was harvested by filtering through 0.45 μm PES filters to remove residual cells and debris. Lentiviruses were either used immediately or aliquoted and stored at −80°C.

### Generation of Stable Cell Lines

hTERT-RPE1 H2B-RFP cells were used to generate cell lines that stably express UBE2H (WT, AA and DD). These cells were subsequently used to generate cell lines that stably express NEK9 (WT and mutants) or AAMP (WT and mutants) by incubating with lentiviruses in the presence of protamine sulfate (2 µg/mL). Following overnight viral infection, cells were switched to fresh media. At 48 h post-infection, antibiotic selection was initiated. For cell lines transduced with pINDUCER20-based lentiviruses, geneticin (Invitrogen, 10131027) was used at 800 µg/mL for 6–7 days. For cell lines transduced with pHAGE-based lentiviruses, puromycin (Sigma Aldrich, P8833) was used at 10 µg/mL for 3 days. Antibiotic-selected populations of cells were expanded as pooled populations and used for further experiments.

### Generation of UBE2H-AA knock-in cell line using CRISPR–Cas9 gene editing

Two hTERT-RPE1 knock-in clones (A3 and B4) expressing UBE2H-S3AS5A (UBE2H-AA) were generated using CRISPR–Cas9 technology by EditCo (Redwood City, CA, USA) according to the manufacturer’s standard protocol.

Guide RNA Sequence: CACCAUGUCAUCUCCCAGUC

Donor Sequence: GGCCCGTGACAGACGGGCCGAGGAAGGGAGAGAGGCGGCGGCGACACCATGTCAG CTCCCGCTCCGGGCAAGAGGCGGATGGACACGGACGTGGTCAAGCTGTATCCTTC

Successful introduction of the S3A, S5A mutation at the endogenous UBE2H locus was confirmed by genomic PCR and Sanger sequencing. Clones were further assessed by karyotype analysis. Expression of UBE2H-AA was confirmed by Western blotting and functional activity was evaluated by detecting UBE2H∼ubiquitin thioesters.

### Small Interfering RNAs (siRNAs)

Cells were transfected using DharmaFECT 1 Transfection Reagent (Horizon, T-2001-02)

according to manufacturer’s instructions with the following siRNAs:

siGENOME NonTargeting Control siRNA #5 (d-001210-05, Dharmacon);

WDR26 siRNA#1 sense, 5’-CCAGAAUAUUGAAGAGGAAUU-3’,

WDR26 siRNA#1 antisense, 5’-UUCCUCUUCAAUAUUCUGGUU-3’;

WDR26 siRNA#2 sense, 5’-GAACAUAGUACAAGAAGAUUU-3’,

WDR26 siRNA#2 antisense, 5’-AUCUUCUUGUACUAUGUUCUU-3’;

MKLN1 siRNA#1 sense, 5’-GCGAAGAGUUGAUUGAAAAUU-3’,

MKLN1 siRNA#1 antisense, 5’-UUUUCAAUCAACUCUUCGCUU-3’;

MKLN1 siRNA#2 sense, 5’-CGAAAUUGGUGUUUGAUCAUU-3’,

MKLN1 siRNA#2 antisense, 5’-UGAUCAAACACCAAUUUCGUU-3’.

Cells were transfected with siRNAs and incubated for 24 h prior to downstream assays. For experiments involving subsequent compound treatment, cells were replaced with fresh media before the addition of compounds.

### *In vitro* screening assay using *Xenopus* egg extract

*Xenopus laevis* egg extracts were prepared as described previously ^44^. Interphase extracts were driven into mitosis by addition of non-degradable cyclin B (MBP-Δ90) as described ^45^. Then, 50 μM HA-tagged ubiquitin (Boston Biochem, U110) was added into the interphase or mitotic extracts for 30 minutes at 24°C. The extracts were then diluted 3 times with XB buffer (10 mM potassium HEPES pH 7.7, 500 mM KCl, 0.1 mM CaCl_2_, 1 mM MgCl_2_, 0.5% NP40, protease inhibitor tablet and 5mM N-ethylmaleimide (NEM)) and incubated with anti-HA beads or control beads for 2h at 4°C. After incubation, beads were removed by centrifugation. The supernatants were collected and processed for mass spectrometry analysis by Tandem mass tag (TMT) method (described below).

### TMT Mass Spectrometry Sample Preparation

#### For mammalian cells

Around 10 million cells were collected and cell pellets were lysed in urea lysis buffer (8M urea, 200 mM EPPS pH 8.0, protease inhibitor tablets, and phosphatase inhibitor tablets) by passing the suspension 20 times through a 21G syringe needle, then protein concentrations were measured protein concentrations by BCA assay (Thermo Fisher Scientific, 23227). Proteins were reduced with 5 mM tris-2-carboxyethyl-phosphine (TCEP, 77720) for 15 min at room temperature, followed by alkylation with 10 mM iodoacetamide for 30 min at room temperature in the dark; excess reagent was then quenched with 15 mM DTT for 15 min at room temperature. Then 100-150 µg of protein were aliquoted out for downstream processing.

#### For *Xenopus* egg extracts

The collected supernatants started directly with the following steps.

Proteins were subsequently purified by methanol-chloroform precipitation, and the pelleted proteins were dissolved in 100 µl of 200 mM EPPS, pH 8.0. LysC (Wako, 125-05061) was added at a 1:100 enzyme-to-protein ratio, and samples were incubated overnight at room temperature with agitation. Trypsin (Pierce, 90305) was then added at a 1:100 enzyme-to-protein ratio and incubated for an additional 6 h at 37 °C. Tryptic digestion was halted by adding 7ml acetonitrile. TMT isobaric reagents (Thermo Fisher Scientific, 90406) were dissolved in anhydrous ACN to 12.5 mg/mL, and peptides were labeled with individual TMT label at a 3:1 label-to-peptide ratio.

Labeling reactions were incubated for 1 h at room temperature with vortex mixing. The reactions were quenched by adding 2.5 µl 5% hydroxylamine (a final concentration of ∼0.3% (v/v)). Equal amounts of each labeled sample were then pooled across all channels at a 1:1 ratio and dried by vacuum centrifugation. Samples were re-suspended in 1% formic acid (FA) in water and desalted using 50 mg 1 cc SepPak C18 cartridge (Waters, WAT054955) under vacuum. Peptides were eluted with 70% ACN/1% FA and lyophilized to dryness by vacuum centrifugation. The combined peptides were fractionated with basic pH reversed-phase (BPRP) HPLC, collected in a 96-well format and consolidated into 24 fractions, of which every second fraction (12 total) was analyzed. Each fraction was desalted using StageTip, dried by vacuum centrifugation, and reconstituted in 5% ACN/5% FA for LC–MS/MS analysis.

### Liquid chromatography and tandem mass spectrometry (*Xenopus* experiments, TMT-MS3)

Mass spectrometric data were collected on Orbitrap Fusion Lumos instruments coupled to a Proxeon NanoLC-1200 UHPLC. The 100 µm capillary column was packed with 35 cm of Accucore 150 resin (2.6 μm, 150Å; ThermoFisher Scientific) at a flow rate of ∼500 nL/min. The scan sequence began with an MS1 spectrum (Orbitrap analysis, resolution 120,000, 400−1400 Th, automatic gain control (AGC) target 400,000, maximum injection time 50ms). Data were acquired 90 minutes per fraction. MS2 analysis consisted of collision-induced dissociation (CID), quadrupole ion trap analysis, automatic gain control (AGC) 20,000, NCE (normalized collision energy) 35, q-value 0.25, maximum injection time 35ms), and isolation window at 0.7 Th. RTS was enabled and quantitative SPS-MS3 scans (resolution of 50,000; AGC target 2.0x10^5^; collision energy HCD at 55%, max injection time of 250 ms) were processed through Orbiter with a real-time false discovery rate filter implementing a modified linear discriminant analysis.

### Database searching (Xenopus experiments, TMT-MS3)

Spectra were converted to mzXML via MSconvert ^46^. Database searching included all entries from the Xenopus reference database. This database was concatenated with one composed of all protein sequences for that database in reversed order. Searches were performed using a 50-ppm precursor ion tolerance and the product ion tolerance was set to 0.9 Da. These wide mass tolerance windows were chosen to maximize sensitivity in conjunction with Comet searches and linear discriminant analysis ^47,48^. TMT (classic) labels on lysine residues and peptide N-termini +229.1629 Da), as well as carbamidomethylation of cysteine residues (+57.021 Da) were set as static modifications, while oxidation of methionine residues (+15.995 Da) was set as a variable modification.

### Liquid chromatography and tandem mass spectrometry (RPE1 experiments, TMT-hrMS2)

Mass spectrometry data were collected using a Orbitrap Eclipse mass spectrometer (Thermo Fisher Scientific, San Jose, CA) coupled with Neo Vanquish liquid chromatograph. Peptides were separated on a 100 μm inner diameter microcapillary column packed with ∼35cm of Accucore C18 resin (2.6 μm, 150 Å, Thermo Fisher Scientific). For each analysis, we loaded ∼2 μg onto the column. Peptides were separated using a 90 min gradient of 5 to 29% acetonitrile in 0.125% formic acid with a flow rate of 350 nL/min. The scan sequence began with an Orbitrap MS^1^ spectrum with the following parameters: resolution 60K, scan range 350-1350, automatic gain control (AGC) target 100%, maximum injection time “auto,” and centroid spectrum data type. We use a cycle time of 1s for MS^2^ analysis which consisted of HCD high-energy collision dissociation with the following parameters: resolution 50K, AGC 200%, maximum injection time 120ms, isolation window 0.6 Th, normalized collision energy (NCE) 36%, and centroid spectrum data type. Dynamic exclusion was set to automatic. The FAIMS compensation voltages (CV) were -30, -50, and -70V.

### Database searching (RPE1 experiments, TMT-hrMS2)

Spectra were converted to mzXML via MSconvert ^46^. Database searching included all entries from the Xenopus reference database. This database was concatenated with one composed of all protein sequences for that database in reversed order. Searches were performed using a 50-ppm precursor ion tolerance and the product ion tolerance was set to 0.03 Da. These wide mass tolerance windows were chosen to maximize sensitivity in conjunction with Comet searches and linear discriminant analysis ^47,48^. TMTpro labels on lysine residues and peptide N-termini +304.207 Da), as well as carbamidomethylation of cysteine residues (+57.021 Da) were set as static modifications, while oxidation of methionine residues (+15.995 Da) was set as a variable modification.

### TMT data analysis

Following database searching, peptide-spectrum matches (PSMs) were adjusted to a 1% false discovery rate (FDR) ^49,50^. PSM filtering was performed using a linear discriminant analysis, as described previously ^48^ and then assembled further to a final protein-level FDR of 1% ^49^. Proteins were quantified by summing reporter ion counts across all matching PSMs, also as described previously ^51^. Reporter ion intensities were adjusted to correct for the isotopic impurities of the different TMTpro reagents according to manufacturer specifications. The signal-to-noise (S/N) measurements of peptides assigned to each protein were summed and these values were normalized so that the sum of the signal for all proteins in each channel was equivalent to account for equal protein loading. Finally, each protein abundance measurement was scaled, such that the summed signal-to-noise for that protein across all channels equals 100, thereby generating a relative abundance (RA) measurement.

### Detection of ubiquitin-charged E2s in *Xenopus* egg extract

Recombinant radiolabeled human UBE2C and UBE2H (WT and mutants) were generated by in vitro transcription/translation in the presence of ^35^S-methionine (Perkin Elmer, NEG709A500UC) with the T7 Coupled Reticulocyte Lysate System (Promega, L1170). Coding sequences for UBE2C and UBE2H were PCR-amplified with gene-specific primers from hORFeome template plasmids and used a template for the translation reactions. *Xenopus* egg extracts were maintained in interphase or pre-arrested in mitosis, then radiolabeled E2s were added and incubated for 0, 15, 30 and 60 min. Reactions were quenched by adding SDS sample buffer lacking reducing agent DTT to preserve E2∼ubiquitin thioesters, and samples were processed for non-reducing SDS–PAGE. Radiolabeled species were detected by phosphor imaging, acquired by a Typhoon laser scanner (Cytiva) and quantified using ImageQuantTL (Cytiva).

### Detection of ubiquitin-charged E2s in cells

To preserve E2∼ubiquitin thioesters, cells pellets were lysed in 50 mM MES buffer, pH 3.5, containing 150 mM NaCl, 2% Nonidet P40, protease inhibitor tablet, and phosphatase inhibitor tablet. Pellets were dispersed by flicking the tube, incubated on ice for 10 min, flicked again, and incubated for an additional 10 min on ice. The lysates were clarified via centrifugation at 15,000g for 10 min at 4 °C. For non-reducing analysis, extract was mixed with 1× NuPAGE™ LDS Sample Buffer (ThermoFisher Scientific, NP0007) without DTT and was subjected to non-reducing SDS–PAGE. To reduce thioester bonds, samples were treated with 1× NuPAGE™ LDS Sample Buffer containing 200 mM DTT and heated for 10 min at 95°C prior to electrophoresis. Proteins were analyzed by western blotting as described below.

### *In vitro* kinase assay

In vitro kinase assays were performed using human UBE2H (Ubiquigent, 62-0032-100) and human CDK2/Cyclin A (Invitrogen, PV3267). Reactions were assembled in TBS buffer with increasing concentrations of CDK2/Cyclin A (0, 0.125, 0.25, and 0.5 μM) and UBE2H (4 μM) with Energy mix (7.5 mM creatine phosphate, 1 mM ATP, 0.1 mM EGTA, 1 mM MgCl2) in a total reaction volume of 30 μL. Reactions were incubated at 37 °C for 5min and quenched in 4× Phos-tag sample buffer supplemented with DTT. Samples were resolved on SuperSep™ Phos-tag™ gels (Fujifilm Wako) using EDTA-free Tris–glycine–SDS running buffer according to the manufacturer’s instructions. Electrophoresis was performed at 110 V for 1.5–2 h on ice. For efficient transfer, gels were incubated in transfer buffer (25 mM Tris, 192 mM glycine, 10% methanol) supplemented with 10 mM EDTA for 10 min and repeated 3 times, followed by equilibration in EDTA-free transfer buffer and transfer to PVDF or nitrocellulose membranes. Gels were stained with Coomassie Brilliant Blue and imaged.

### Western blotting

Cell pellets were resuspended in 2% SDS buffer (2% SDS, 50 mM Tris-HCl pH 8.0, 10 mM EDTA). Pellets were dispersed by flicking and lysates were homogenized by repeated pipetting and/or brief vortex until cells were fully lysed. Samples were heated at 95°C for 10 min, cooled to room temperature, and briefly spun down. To reduce viscosity and improve uniformity, lysates were sonicated for 10 s. Protein concentrations were determined by BCA assay and normalized lysate amounts were mixed with 1× NuPAGE™ LDS Sample Buffer (ThermoFisher Scientific, NP0007) and heated prior to electrophoresis. Proteins were separated on NuPAGE 4–12% Bis-Tris gel (ThermoFisher Scientific) and transferred via wet transfer onto PVDF membranes (ThermoFisher Scientific, 88518). Membranes were probed with the indicated primary antibodies and corresponding HRP- or fluorescent dye–conjugated secondary antibodies. Chemiluminescent blots were developed using Chemiluminescent Substrate (ThermoFisher Scientific, 34580) and imaged using Amersham™ ImageQuant™ 800 (Cytiva). For fluorescence-based detection, immunoblots were developed using LI-COR secondary antibodies and imaged using Odyssey CLx Imager (LICORbio). Band intensities were quantified using ImageQuantTL (Cytiva) and normalized to the indicated loading control (Vinculin or GAPDH).

### Time Lapse and Fluorescence Microscopy

For the live-cell imaging experiments tracking chromosome segregation and mitotic duration, hTERT-RPE1 H2B-RFP cell lines that stably expressing UBE2H (WT, AA and DD) and UBE2H knock-in cells were used. Cells were plated in 12- or 24-well glass-bottom plates (Greiner BioOne, 662892). After 24 h, cells were treated with the indicated compounds (e.g. nocodazole). For hTERT-RPE1 H2B-RFP cell lines, time-lapse imaging was initiated immediately. For UBE2H knock-in cells, siR-Hoechst (200 nM) was added to cells and incubated for 4 h at 37°C before performing time-lapse imaging. Imaging was performed on an inverted fluorescence microscope (Nikon Eclipse Ti-E with Perfect Focus system) equipped with a motorized stage (Prior ProScan III), a 20× air objective, using a stage-top incubator (Okolab) maintained at 37°C with 5% CO_2_ with humidification. Differential interference-contrast and fluorescence images (mCherry or SiR-Hoechst channels) were acquired from two fields of view per condition at 3-10-min intervals for 24–48 h, depending on the experiment. Fluorescence was excited with a Lumencor Spectra-X using a 555/25 excitation filter and a 605/52 emission filter (Chroma). siR-Hoechst fluorescence was excited using a 640/30 excitation filter set (ET-Cy5 narrow filter set, 49009, Chroma) and emission detection around ∼690 nm. Videos were analyzed using Nikon NIS-Elements AR. Mitotic duration was defined as the time from nuclear envelope breakdown (NEB) to cell division, death (cytoplasmic blebbing), or mitotic slippage.

### Growth curve analysis

Viable cell numbers were quantitated at the indicated timepoint using CellTiter-Glo® Cell Viability Assay (Promega) and luminescence was recorded with the Synergy 2 microplate reader (BioTek). For each condition, values were normalized to the baseline measurement to calculate fold change in viable cell number over time. For growth curve comparison across conditions, cells were seeded at equal starting densities and growth values were normalized to the day-0 measurement.

### Cell-cycle analysis by propidium iodide (PI) staining

Cells were harvested and washed with PBS, then fixed by dropwise addition of cold 70% ethanol while vortexing and incubated at −20°C (≥2 h or overnight). Fixed cells were washed to remove ethanol and incubated in PI staining solution containing propidium iodide (50 µg/mL) and RNase A (100 µg/mL) in PBS for 30 min at room temperature protected from light. DNA-content profiles were acquired with an Attune NxT Flow Cytometer. The fractions of cells in G1, S and G2/M were quantified using FlowJo.

### Colony formation assay

Cells were seeded at low density in 6-well plates (500–1,000 cells per well) in complete growth medium and allowed to attach overnight. After 14 days, cells were washed with PBS, fixed with 4% paraformaldehyde for 15 min at room temperature, and stained with 0.5% crystal violet in 20% methanol for 30 min. Excess stain was removed by rinsing with water and plates were air-dried. Colonies were imaged, and colony area was quantified in ImageJ by thresholding images followed by particle detection using the Analyze Particles function with an area cutoff to exclude small artifacts.

### IP-MS to identify AAMP interactions

To identify AAMP-associated proteins, cells stably expressing N-terminally HA-tagged wild-type AAMP or two stable AAMP mutants (△C429-434 and D433A) were harvested and lysed for 20 min at 4°C in TBS buffer (30 mM Tris-HCl pH 7.4, 150 mM NaCl) containing 0.5% NP-40, protease inhibitor tablet, and phosphatase inhibitor tablet. Lysates were clarified by centrifugation at 10,000g for 10 min, and cleared supernatants were incubated with anti-HA beads for 4h at 4°C. The beads were washed three times with wash buffer (TBS containing 0.1% NP-40) and bound proteins were eluted twice using 250 μg/mL HA peptide in PBS at 37°C for 30-minute incubations with gentle shaking. Purified samples were precipitated by chloroform–methanol and digested using Pierce Trypsin (Pierce, 90305, 1 mg/mL stock) at a 1:100 enzyme-to-protein ratio for 6 h at 37°C, then processed for label-free LC–MS/MS proteomic analysis and quantified the number of peptide spectral counts in each group.

### Mass spectrometric data collection (IP-MS)

Mass spectrometry data were collected using a Exploris 480 mass spectrometer (Thermo Fisher Scientific, San Jose, CA) coupled with a Proxeon 1200 Liquid Chromatograph (Thermo Fisher Scientific). Peptides were separated on a 100 μm inner diameter microcapillary column packed with ∼25 cm of Accucore C18 resin (2.6 μm, 150 Å, Thermo Fisher Scientific). We loaded ∼0.5 μg onto the column.

Peptides were separated using a 60min gradient of 3 to 22% acetonitrile in 0.125% formic acid with a flow rate of 350 nL/min. The scan sequence began with an Orbitrap MS^1^ spectrum with the following parameters: resolution 60,000, scan range 350−1350 Th, automatic gain control (AGC) target “standard”, maximum injection time was set to 50ms, RF lens setting 50%, and centroid spectrum data type. We selected the top twenty precursors for MS^2^ analysis which consisted of HCD high-energy collision dissociation with the following parameters: resolution 15,000, AGC was set at “standard”, maximum injection time “auto”, isolation window 1.2 Th, normalized collision energy (NCE) 28, and centroid spectrum data type. In addition, unassigned and singly charged species were excluded from MS^2^ analysis and dynamic exclusion was set to 90 s. Each sample was run twice, once with CV set of -30/-50/-70 V and again with a CV set of - 40/-60/-80V. A 1 s TopSpeed cycle was used for each CV.

### Structural analysis

Structural predictions were performed using AlphaFold 3 (https://alphafoldserver.com/), as described ^35^, and models were visualized using UCSF ChimeraX version 1.8rc202405310011 (2024-05-31) https://www.rbvi.ucsf.edu/chimerax. ipTM and pTM scores for interaction predictions are provided in Supplementary Table 6.

### Quantification and statistical analysis

Proteomics datasets were processed and analyzed in R Project (R Foundation for statistical computing) using custom scripts. Data visualizations in the form of volcano plots were generated using the online tool EasyPubPlot. All statistical data were calculated using GraphPad Prism 10 by the built-in one-way repeated-measures ANOVA with Dunnett multiple-comparisons tests against the pre-specified control condition. A significance threshold of p < 0.05 was applied. No correction for multiple comparisons was applied, and reported p-values are unadjusted. The number of independent experimental repeats is reported in the corresponding figure legends.

## Acknowledgements

This work was supported by NIH grants R01 GM132129 (J.A.P.), R01 GM67945 (S.P.G.), and R35 GM127032 (R.W.K.). We thank D. Finley, B. Manning, A. Sharma and the other members of the KLHF groups for helpful discussions. We thank J.W. Harper for the gift of plasmids (hORFeome collection, pPAX2, pMD2 and pHAGE-FLAG-HA-NTAP vector) and T. Fu and C. Lee for their helpful suggestions regarding plasmid construction and lentivirus-based experiments. We thank A. Holland for the gift of hTERT-RPE1 cells stably expressing H2B-RFP. We thank S. Elledge for the gift of pINDUCER20. We thank B. Manning for sharing the TSC2 knockout clones and J. Shin for his suggestions on working with these clones. We thank for the gift of PFI-7 from C. H. Arrowsmith of the Structural Genomics Consortium. We thank J. Waters and the Core for Imaging Technology & Education at Harvard Medical School for technical assistance. We thank A. Dandapat and W. Trone for their dedicated contributions during their summer internships. The mass spectrometry proteomics data have been deposited in the ProteomeXchange Consortium.

## Author contributions

Y.C. designed, performed, and analyzed all proteomic and cell biological experiments unless otherwise mentioned. V.R. designed and analyzed the *in vitro* proteomic screen in *Xenopus* egg extract with assistance from L.F. and J.A.P; V.R. performed all *Xenopus* egg extract experiments and made the original observation about UBE2H mitotic-dependent regulation; V.R. and S.M. performed the site-directed mutagenesis and cloning to generate the AA and DD mutants and made the initial observations of ubiquitin charging phenotypes in these mutants. J.A.P. and S.P.G. performed TMT-mass spectrometric analysis; M.K. performed live-cell imaging experiments in UBE2H-inducing cells; S.M. detected endogenous UBE2H thioester in RO3306-arrested and release cells; N.O. performed *in vitro* phosphorylation experiment. R.W.K. conceived and supervised this study and assisted with experimental design and interpretation; Y.C. and R.W.K. wrote the manuscript with input from other authors.

---

**Extended Data Figure 1.**
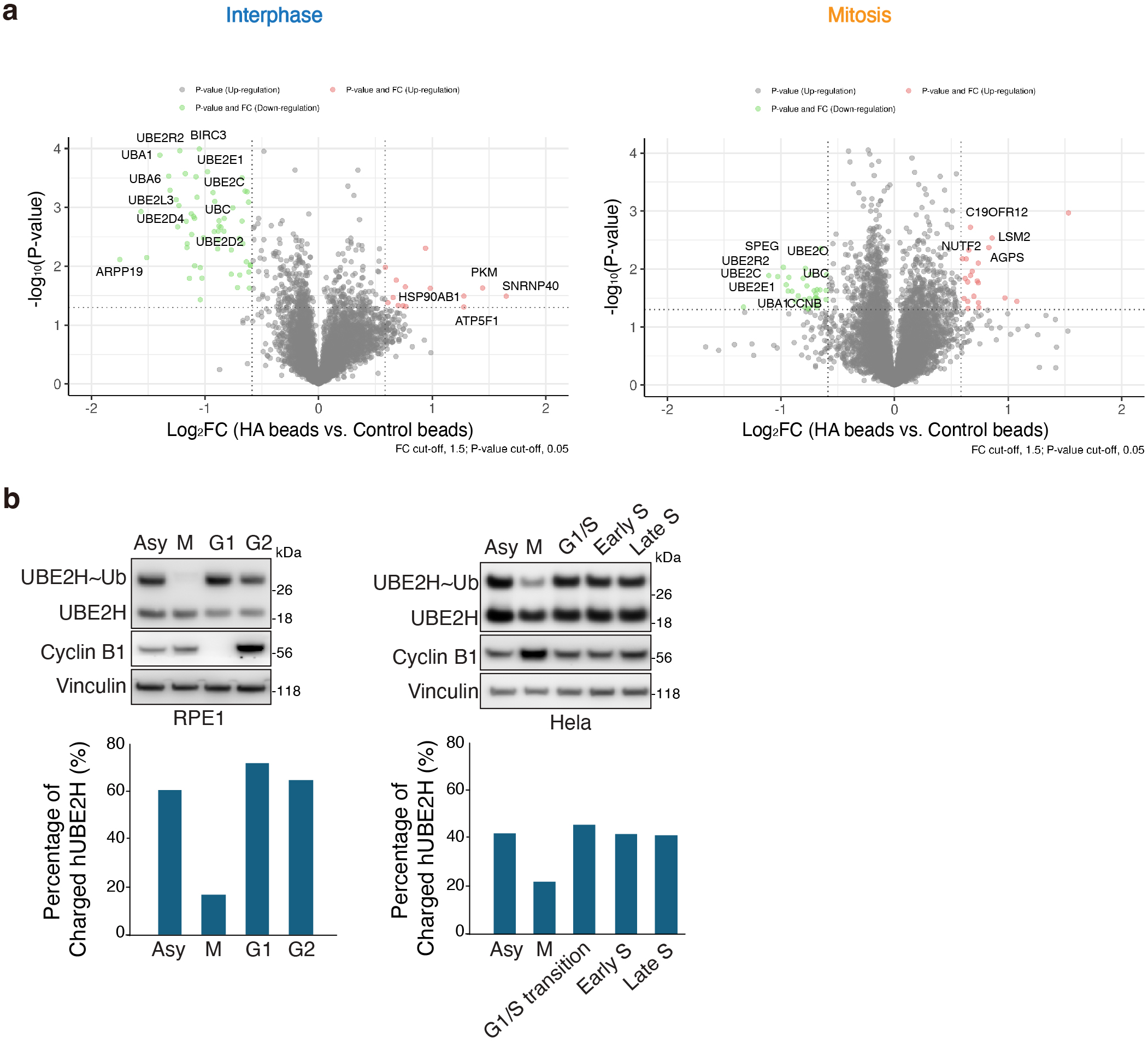
**a.** Volcano plots of -log10 p-value versus the log2 fold change of anti-HA beads vs control beads in Xenopus egg extracts in interphase and mitosis. Two or three biological replicates were analyzed for each of four conditions. In total, 7,044 proteins were quantified across all conditions. The downregulated proteins are indicated in green, and the upregulated proteins are indicated in red. Top regulated proteins are labeled. **b.** Levels of ubiquitin-charged endogenous UBE2H in hTERT-RPE1and Hela cells. For hTERT-RPE1 cells, mitotic arrest was induced by nocodazole (600 nM) for 18h; G1 arrest with palbociclib (1 µM) for 20h; G2 arrest with RO-3306 (7.5 µM) for 18h. For Hela cells, mitotic arrest was induced with nocodazole (600 nM) for 18h; G1/S arrest by a double-thymidine block (2 mM thymidine overnight, release for 8 h, followed by a second block with 2 mM thymidine overnight); early S phase, 2 h after release; late S phase, 4 h after release. Cell lysates were analyzed under non-reducing conditions to examine UBE2H∼Ub (top). Quantification of the percentage of ubiquitin-charged E2s is shown (bottom).

**Extended Data Fig 2.**
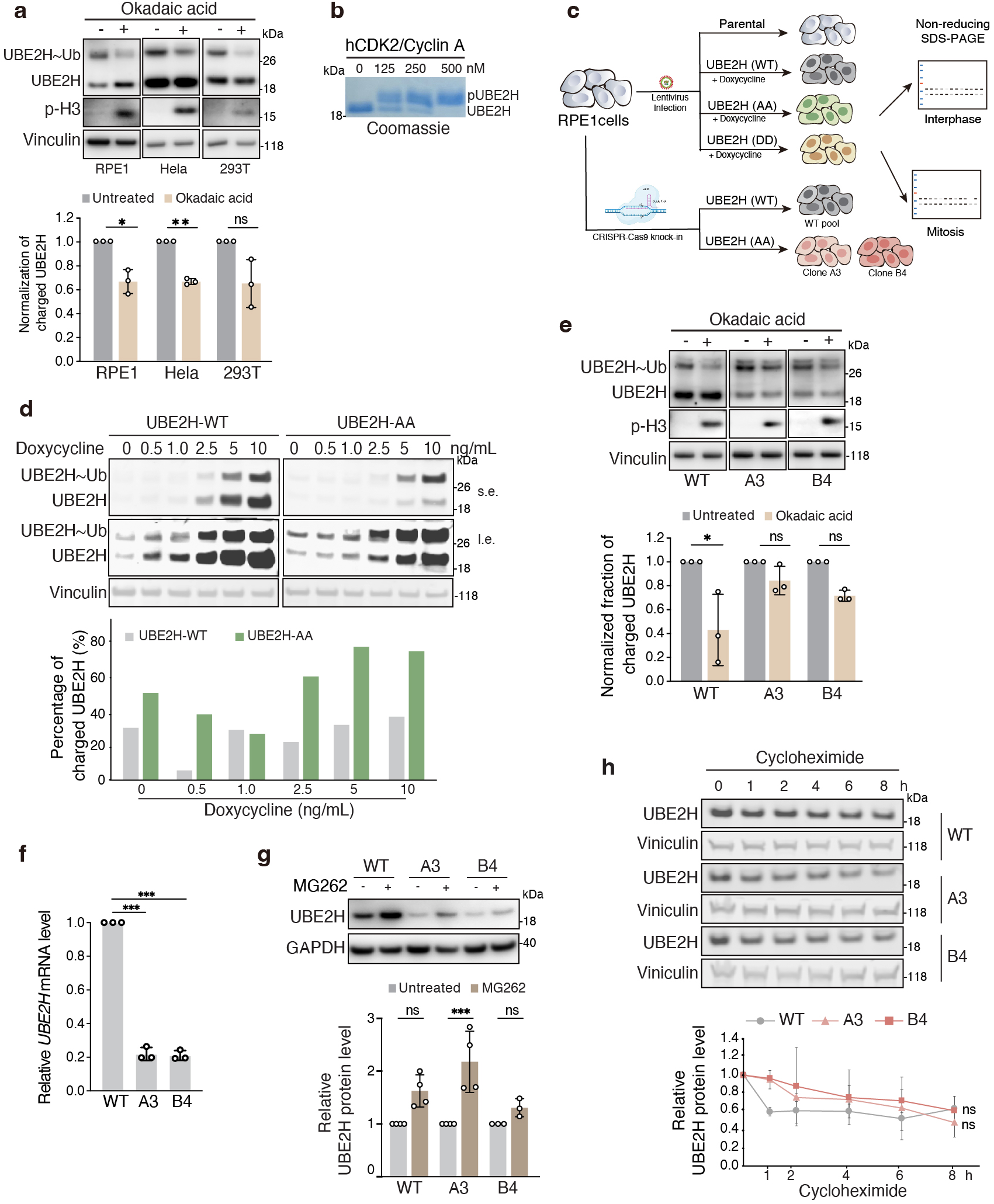
**a.** Levels of ubiquitin-charged endogenous UBE2H in hTERT-RPE1, Hela and HEK293T cells_ Cells were treated with okadaic acid (200 nM) for 3h_ Cell lysates were analyzed under non-reducing conditions to examine levels of UBE2H∼Ub (top)_ Quantification represents mean± s_d_ from three independent biological replicates_ **b.** Phosphorylation of purified UBE2H in the presence of increasing concentrations of recombinant CDK2/Cyclin A was measured using a Phos-tag gel assay_ **c.** Schematic of the generation of stable hTERT-RPE1 cells expressing doxycycline-inducible UBE2H constructs (WT, AA and DD) by lentiviral transduction and application of CRISPR-Cas9 editing of the endogenous UBE2H locus to introduce the AA mutation (two independent clones, A3 and B4)_ **d.** Levels of doxycycline-inducible ubiquitin-charged UBE2H (WT, AA) were examined by anti-UBE2H immunoblotting in stably expressing hTERT-RPE1 cells_ Cells were induced by increasing concentrations of doxycycline (0, 0.5, 1, 2.5, 5 and 1Ong/ml)_ s.e., short exposure; Le., long exposure_ Quantification of the percentage of ubiquitin-charged UBE2H is shown (bottom)_ **e.** Levels of ubiquitin-charged endogenous UBE2H in WT cells and knock-in clones_ Cells were treated with okadaic acid (200 nM) for 3h_ Cell lysates were analyzed under non-reducing conditions to examine levels of UBE2H∼Ub (top)_ Quantification represents mean± s_d_ from three independent biological replicates (bottom)_ **f.** Relative UBE2H mRNA levels in WT cells and knock-in clones_ Quantification represents mean± s_d_ from three independent biological replicates_ **g.** UBE2H protein levels in WT cells and knock-in clones with or without the proteasome inhibitor MG-262_ Quantification of relative UBE2H protein level represents mean ± s_d_ from three independent biological replicates_ **h.** UBE2H degradation in WT cells and knock-in clones in the presence of cycloheximide (CHX) as measured by immunoblotting_ Quantification of relative UBE2H protein level represents mean± s_d_ from three independent biological replicates_ Statistical significance of differences between groups was determined with one-way ANOVA analysis **(f)** or two-tailed unpaired Student’s t-test **(a, e, g-h)_** *** = p < 0.001 _ ** = p < 0.01 _ * = p < 0.05_ ns = not statistically significant

**Extended Data Fig. 3.**
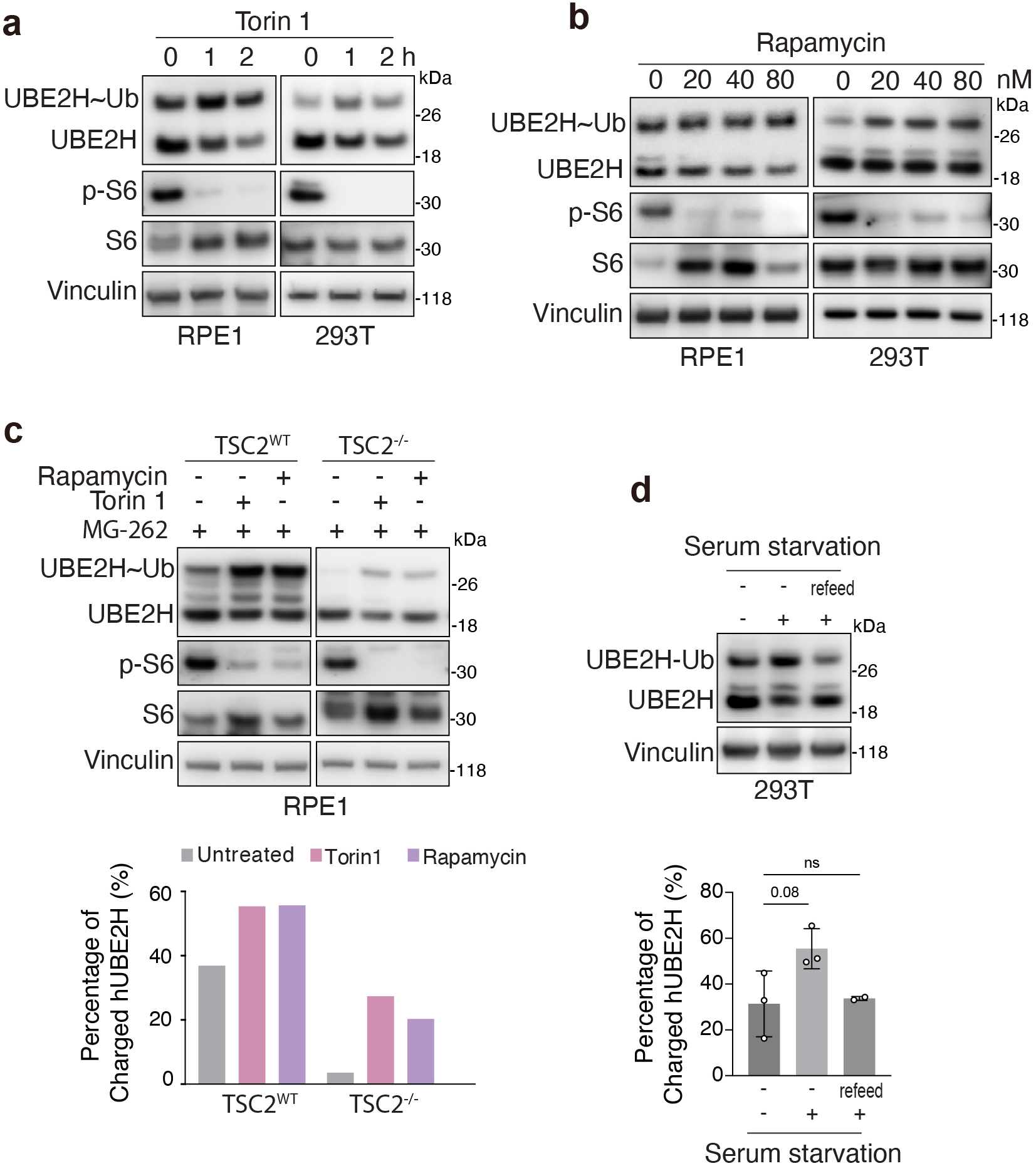
**a.** Non-reducing SOS-PAGE analysis of UBE2H thioester-linked ubiquitin conjugates in hTERT-RPE1and HEK293T cells after treatment with torin1 (250 nM) for the indicated time. Quantification represents mean± s.d. from three independent biological replicates. **b.** Non-reducing SOS-PAGE analysis of UBE2H thioester-linked ubiquitin conjugates in hTERT-RPE1 cells and HEK293T cells after treatment with rapamycin at the indicated concentrations for 1 h. Quantification of the percentage of ubiquitin-charged UBE2H is shown (bottom). **c.** Non-reducing SOS-PAGE analysis of UBE2H thioester-linked ubiquitin conjugates after treatment with torin1 (250 nM) and rapamycin (80 nM) in WT, *Tsc2-^1^-* hTERT-RPE1cells. Quantification of the percentage of charged UBE2H is shown (bottom). **d.** Non-reducing SOS-PAGE analysis of UBE2H thioester-linked ubiquitin conjugates in HEK293T cells that were incubated in serum-depleted medium for 12 h. Quantification represents mean ± s.d. from three independent biological replicates. Statistical significance of differences between groups was determined with one-way ANOVA analysis. * = p < 0.05. ns = not statistically significant.

**Extended Data Fig. 4.**
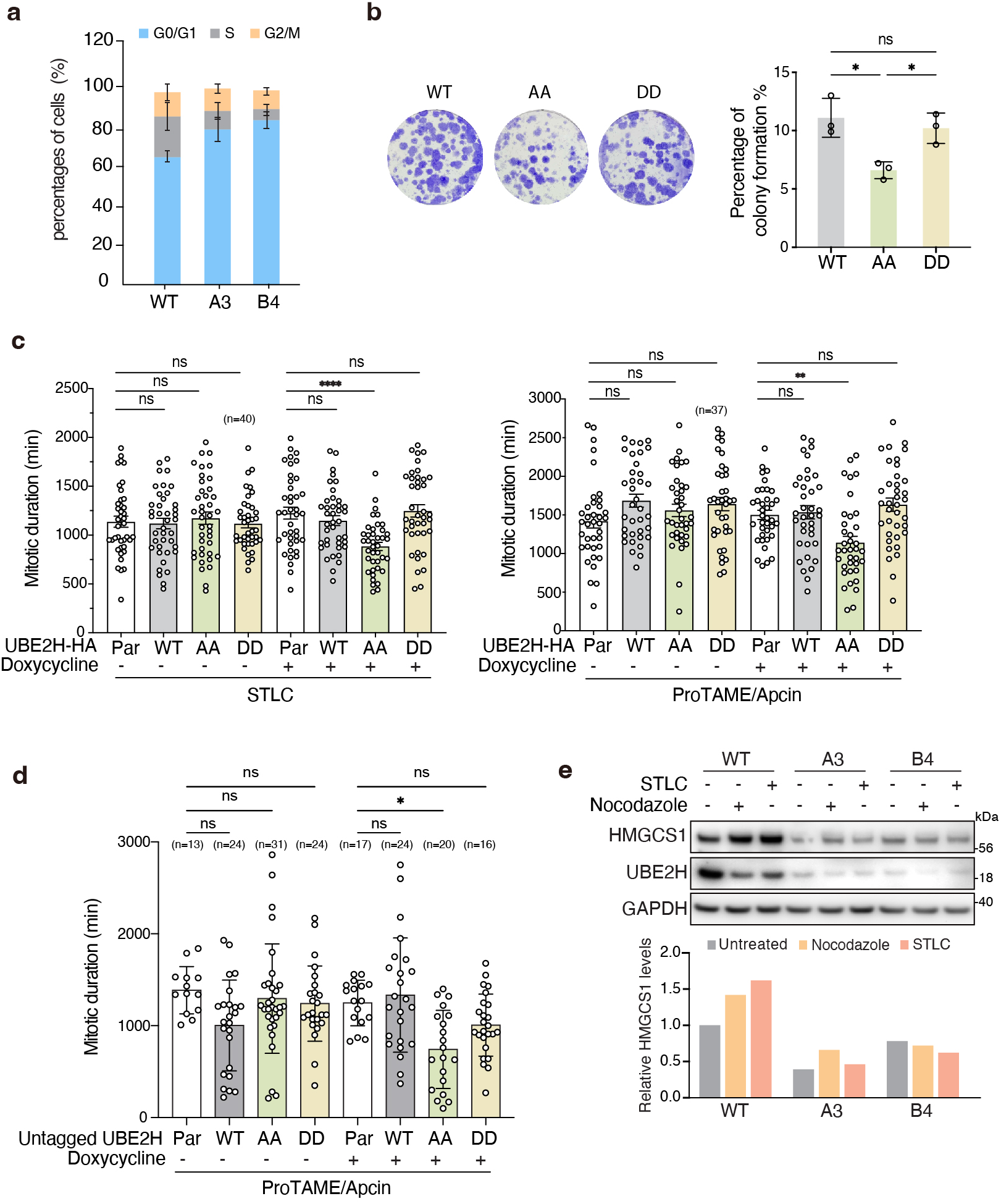
**a.** FAGS analysis of propidium iodide (Pl)-stained WT cells and knock-in clones (A3 and B4). Percentages of cells in each cell cycle phase were calculated. Quantification represents mean ± s.d. from three independent biological replicates. **b.** Colony formation assays in the hTERT-RPE1 cells that stably expressing doxycycline-inducible UBE2H constructs (WT, AA and DD) treated with doxycycline (5 ng/ml) for 14 days. Quantification represents mean ± s.d. from three independent biological replicates. **c.** Mitotic arrest duration of hTERT-RPE1 cells that stably express doxycycline-inducible UBE2H constructs (WT, AA and DD). Cells were treated with STLC (1.5 µM) or proTAME (12.5 µM) and apcin (25 µM) and imaged every 1O mins by wide-field time-lapse microscopy for 48 h. Quantification was performed on the number of cells indicated on the plot. Each point represents an individual cell’s mitotic duration, measured as the time from NEB to division, slippage, or cell death. **d.** Mitotic arrest duration of hTERT-RPE1 cells that stably express doxycycline-inducible untagged UBE2H constructs (WT, AA and DD). Cells were treated with proTAME (12.5 µM) and apcin (25 µM) and imaged every 1O mins by wide-field time-lapse microscopy for 48 h. Quantification was performed on the number of cells indicated on the plot. Each point represents an individual cell’s mitotic duration, measured as the time from NEB to division, slippage, or cell death. **e.** Western blots showing HMGCS1, UBE2H protein levels in asynchronous WT cells and knock-in clones. Mitotic cells were treated with nocodazole (600 nM) or STLC (2 µM) for 18h and collected by shake-off. Quantification of the relative HMGCS1 protein levels is shown (bottom). For b-c, Error bars= mean±s.d. p-values were calculated by one-way ANOVA. **** = p < 0.0001. ** = p < 0.01. * = p < 0.05. ns = not statistically significant.

**Extended Data Fig. 5.**
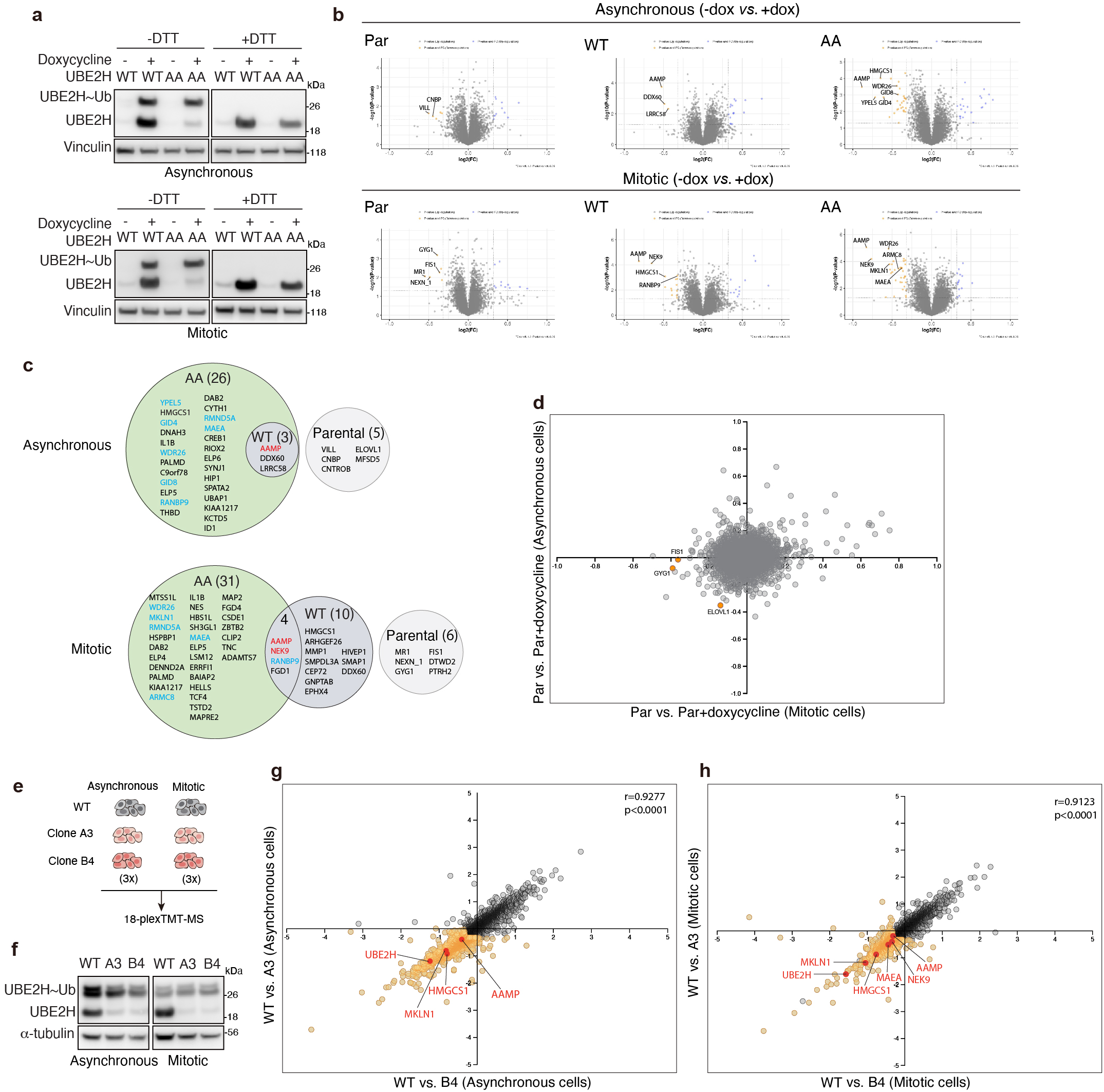
**a.** Western blotting of cells used for the proteomics experiment. Levels of doxycycline-inducible ubiquitin-charged UBE2H (WT, AA) were examined by anti-UBE2H immunoblotting in stable hTERT-RPE1 cells with or without the presence of OTT. **b.** Volcano plots of whole cell proteomic analysis of parental cells and UBE2H (WT, AA) cells treated with doxycycline compared with untreated cells in asynchronous and mitotic conditions. Shown are -109_10_ (P-value) (y-axis) vs Log_2_ (fold change in protein abundance) (x-axis) for doxycycline-treated versus untreated cells, from three independent biological replicates. **c.** Venn diagrams showing overlap among proteins downregulated by doxycycline addition in parental cells or those overexpressing UBE2H (WT, AA) (FC < -1.25, p <0.05 and number of peptides >1), in asynchronous and mitotic conditions. **d.** Two-dimensional plots comparing proteins downregulated in parental cells after treatment with doxycycline in asynchronous and mitotic conditions. Significantly downregulated proteins are indicated in orange. **e.** Schematic of the TMT-mass spectrometry workflow to identify proteins differentially expressed in asynchronous and mitotic conditions in WT cells and knock-in clones across three independent replicates. **f.** Levels of ubiquitin-charged UBE2H were examined by anti-UBE2H immunoblotting in WT cells and knock-in clones in asynchronous and mitotic conditions. **g-h.** Two-dimensional plots of TMT-MS-based quantitation of protein abundance in the two knock-in clones under asynchronous and mitotic conditions. Each dot represents a quantified protein; axes indicate the log2-fold change in A3 (y-axis) and B4 (x-axis) relative to WT. Pearson’s correlation analysis revealed a strong positive concordance between the two clones (asynchronous: r = 0.9277, p < 0.0001; mitotic: r = 0.9123, p < 0.0001). HMGCS1, MKLN1, UBE2H, AAMP, MAEA and NEK9 are highlighted in red, and the other downregulated proteins are indicated in orange.

**Extended Data Fig. 6.**
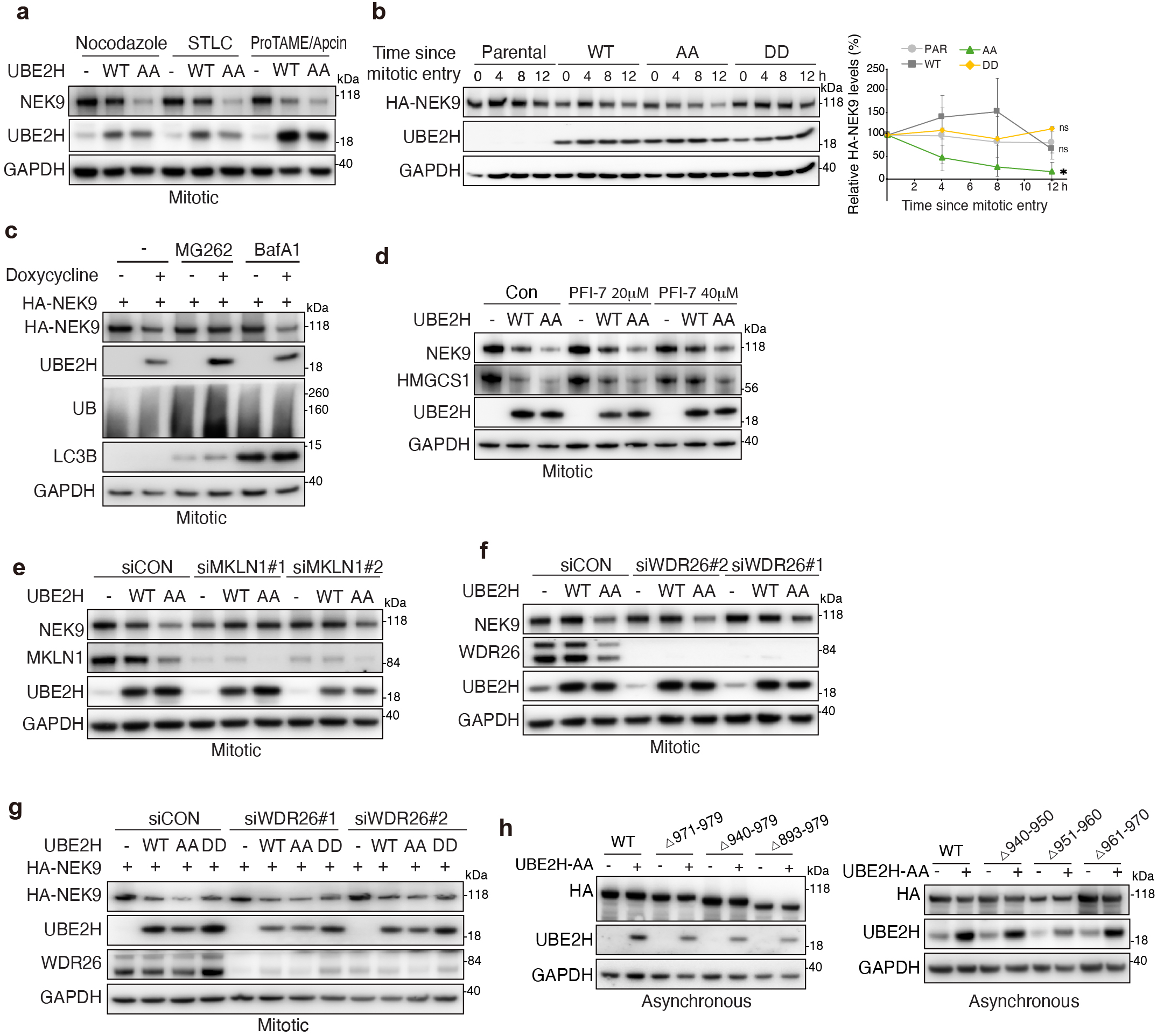
**a.** Western blots showing NEK9 protein levels in mitotic parental and UBE2H (WT, AA) hTERT-RPE1 cells after treatment with nocodazole (600 nM), STLC (2 µM), proTAME (12. 5 µM)/ apcin (25 µM). **b.** lmmunoblots of NEK9 in hTERT-RPE1 cells. Cells were arrested by RO3306 7.5 µM for 18h, then released into nocodazole for the indicated time, and mitotic cells were harvested and analyzed by immunoblotting. Quantification represents mean± s.d. from three independent biological replicates. **c.** Western blots showing exogenous NEK9 protein levels in mitotic UBE2H-AA-expressing hTERT-RPE1 cells after treatment with MG262 (10 µM) or BafA1 (1 µM) in the presence or absence of doxycycline. **d.** Western blots showing NEK9 protein levels in mitotic parental and UBE2H (WT, AA) hTERT-RPE1 cells after treatment with PFl-7 (40 µM). **e.** Western blots showing NEK9 protein levels in mitotic parental and UBE2H (WT, AA) hTERT-RPE1 cells transfected with siCON or MKLN1 siRNAs. **f.** Western blots showing NEK9 protein levels in mitotic parental and UBE2H (WT, AA) hTERT-RPE1 cells transfected with control or WDR26 siRNAs. **g.** Western blots showing exogenous NEK9 protein levels in mitotic parental and UBE2H (WT, AA, DD) hTERT-RPE1 cells transfected with siCON or WDR26 siRNAs. **h.** Western blots showing levels of NEK9 WT or truncation mutants in asynchronous UBE2H-AA-expressing hTERT-RPE1 cells in the presence or absence of doxycycline. Statistical significance of differences between groups was determined with two-tailed unpaired Student’s t-test (b). * = p < 0.05. ns = not statistically significant.

**Extended Data Fig. 7.**
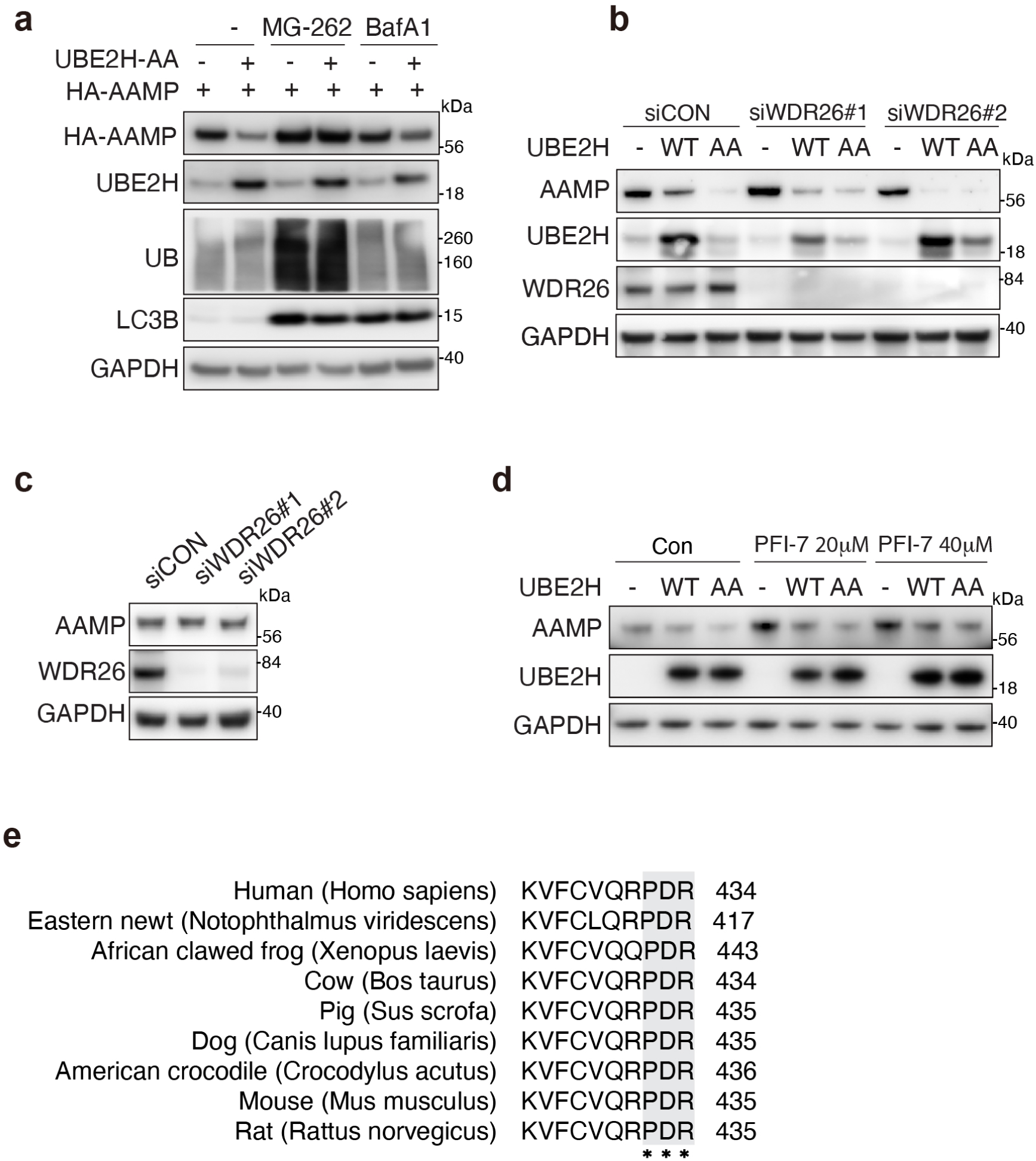
**a.** Western blots showing exogenous AAMP protein levels in UBE2H-AA-expressing hTERT-RPE1 cells after treatment with MG-262 (10 µM) or BafA1 (1 µM) in the presence or absence of doxycycline. **b.** Western blots showing AAMP protein levels in parental and UBE2H (WT, AA) hTERT-RPE1 cells transfected with siCON or WDR26 siRNAs. **c.** Western blots showing AAMP protein levels in hTERT-RPE1 cells transfected with siCON or WDR26 siRNAs. **d.** Western blots showing AAMP protein levels in parental and UBE2H (WT, AA) hTERT-RPE1 cells after treatment with PFl-7. **e.** Multiple sequence alignment of the indicated protein C-terminal region from representative vertebrate species. The conserved PDR motif is highlighted (boxed). Residue numbering corresponds to each sequence.

**Extended Data Fig. 8.**
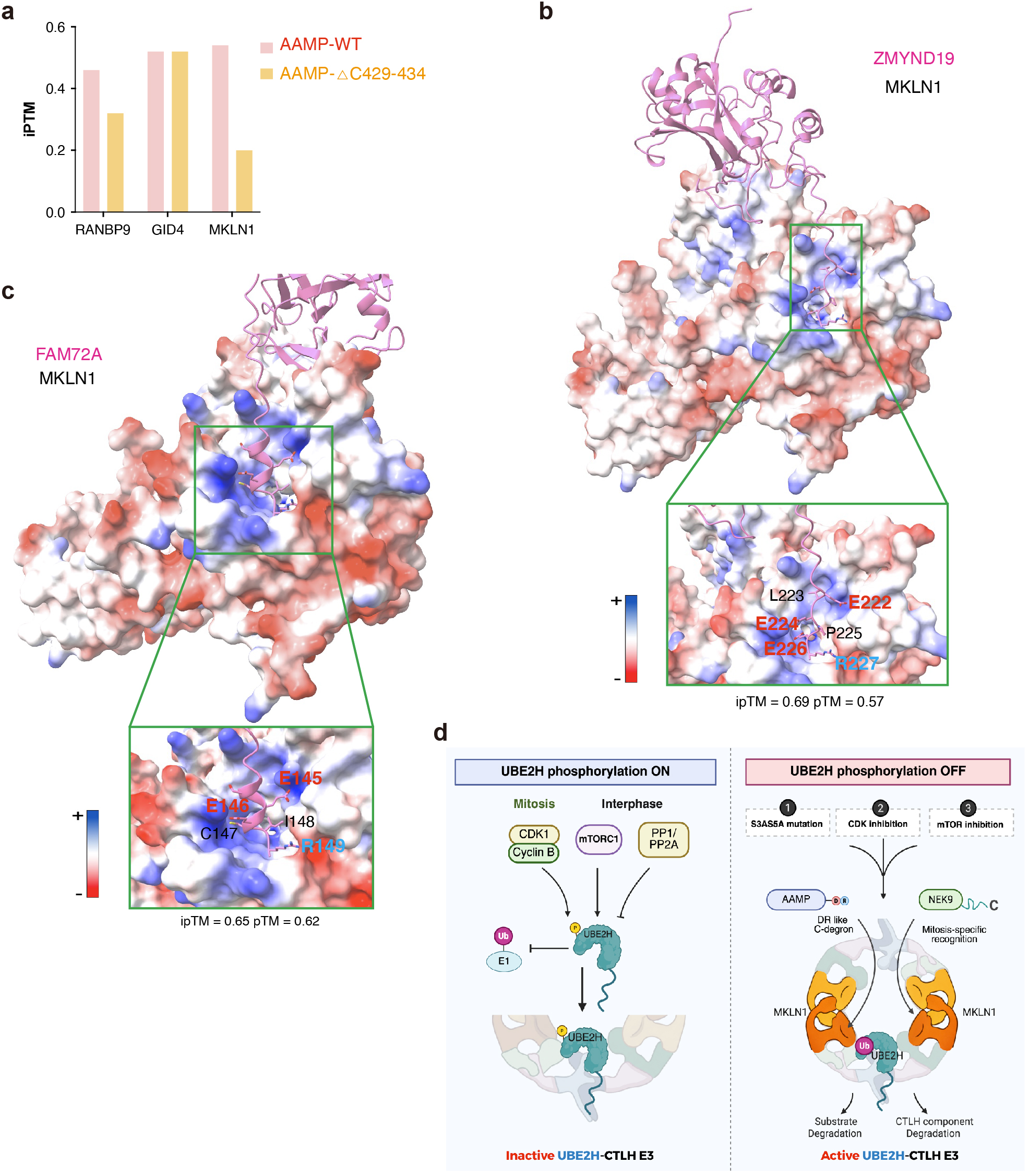
**a.** Predicted interactions between AAMP (WT, ΔC429-434) and CTLH components (RANBP9, GID4, MKLN1) using AlphaFold 3, showing ipTM scores. **b.** AF3 models for ZMYND19 (pink) bound to MKLN1, shown as an electrostatic potential surface. Residues at the extreme ZMYND19 C-terminus interact with a groove formed by the blades of MKLN1 β-propeller. ipTM scores for pairwise combinations are indicated. **c.** AF3 models for FAM72A (pink) and MKLN1 (electrostatic potential surface). Residues at the FAM72A C-terminus interact with the groove formed by blades of the MKLN1 β-propeller. ipTM scores for pairwise combinations are indicated. **d.** Graphical summary. Mitotic CDK- and interphase mTOR-dependent phosphorylation of UBE2H at N-terminal Ser3/Ser5 reduces UBE2H∼Ub thioester formation ("charging"), thereby limiting the pool of active E2 available to CTLH. Preventing S3/S5 phosphorylation maintains UBE2H charging enhances CTLH-mediated substrate degradation (e.g. NEK9 and AAMP), and a DR-like C-terminal degron recognized by the CTLH subunit MKLN1 is defined.

