## Extended Data Figures for "CDK/mTOR–dependent phosphorylation of UBE2H restrains its charging with ubiquitin and regulates CTLH-dependent degradation"

### Extended Data Figure 1

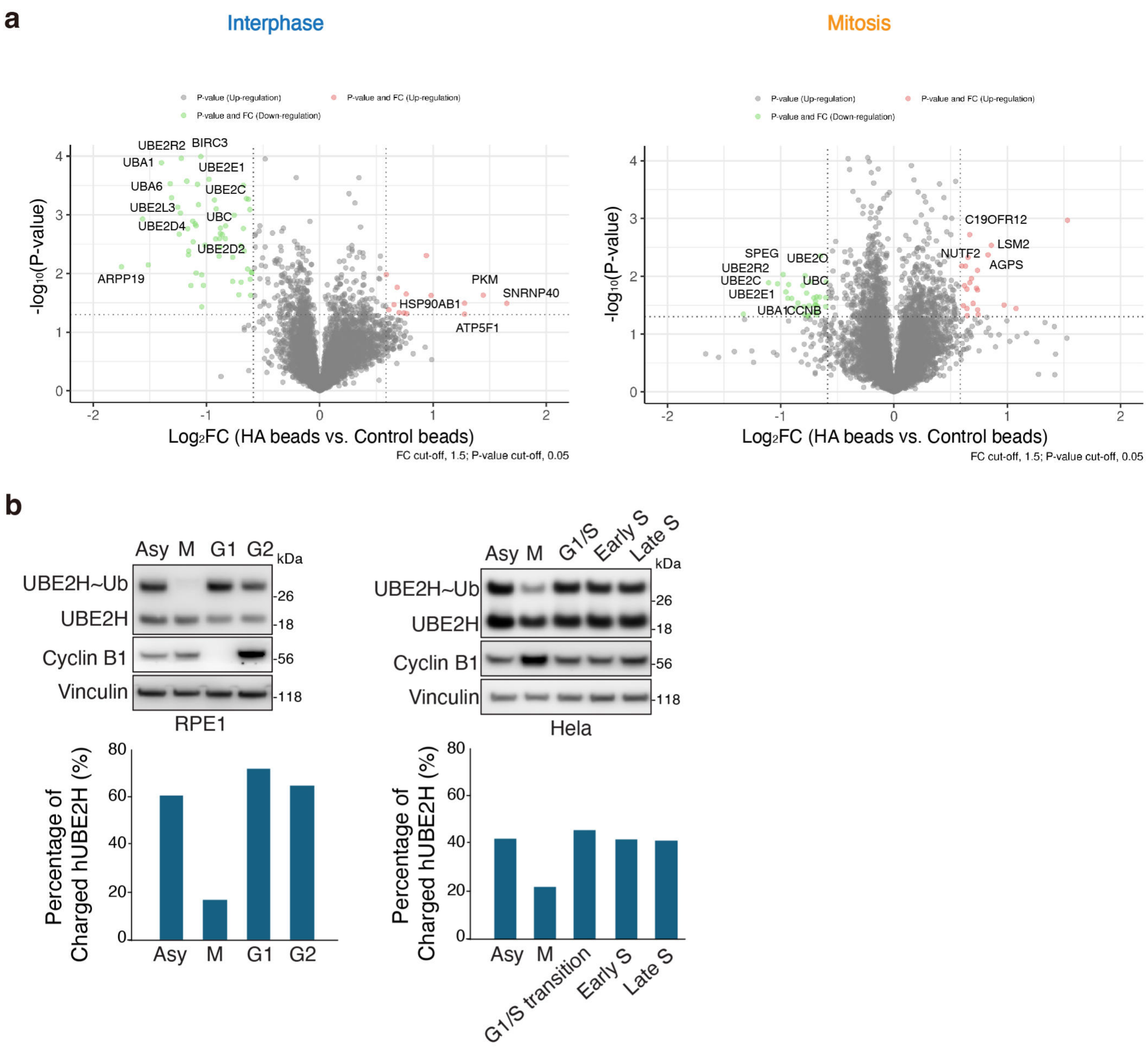


**Extended Data Figure 1. a.** Volcano plots of –log10 p-value versus the log2 fold change of anti-HA beads vs control beads in Xenopus egg extracts in interphase and mitosis. Two or three biological replicates were analyzed for each of four conditions. In total, 7,044 proteins were quantified across all conditions. The downregulated proteins are indicated in green, and the upregulated proteins are indicated in red. Top regulated proteins are labeled. **b.** Levels of ubiquitin-charged endogenous UBE2H in hTERT-RPE1and HeLa cells. For hTERT-RPE1 cells, mitotic arrest was induced by nocodazole (600 nM) for 18h; G1 arrest with palbociclib (1 µM) for 20h; G2 arrest with RO-3306 (7.5 µM) for 18h. For HeLa cells, mitotic arrest was induced with nocodazole (600 nM) for 18h; G1/S arrest by a double-thymidine block (2 mM thymidine overnight, release for 8 h, followed by a second block with 2 mM thymidine overnight); early S phase, 2 h after release; late S phase, 4 h after release. Cell lysates were analyzed under non-reducing conditions to examine UBE2H~Ub (top). Quantification of the percentage of ubiquitin-charged E2s is shown (bottom).

### Extended Data Figure 2

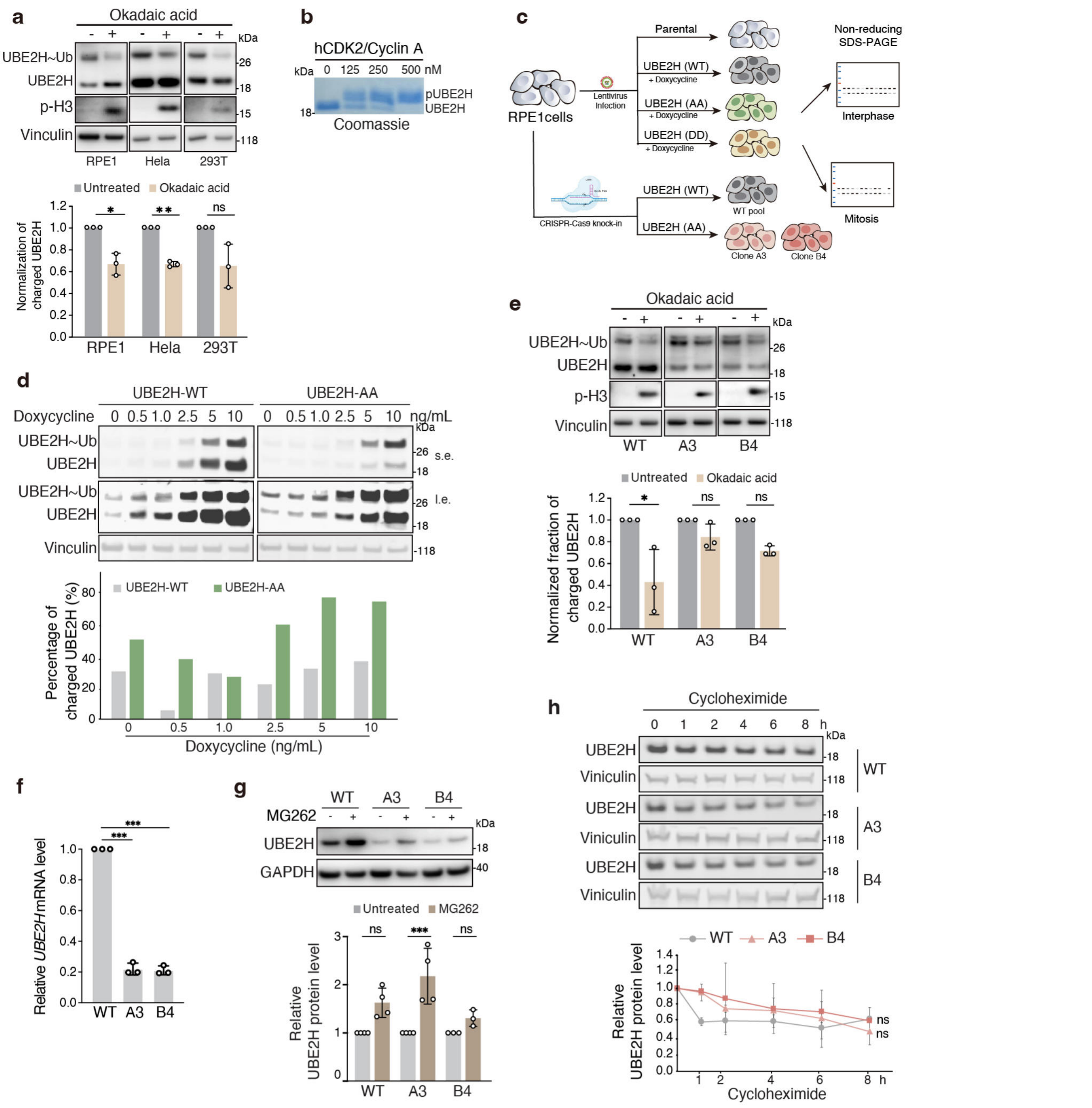


**Extended Data Fig 2. a.** Levels of ubiquitin-charged endogenous UBE2H in hTERT-RPE1, HeLa and HEK293T cells. Cells were treated with okadaic acid (200 nM) for 3h. Cell lysates were analyzed under non-reducing conditions to examine levels of UBE2H~Ub (top). Quantification represents mean ± s.d. from three independent biological replicates. **b.** Phosphorylation of purified UBE2H in the presence of increasing concentrations of recombinant CDK2/Cyclin A was measured using a Phos-tag gel assay. **c.** Schematic of the generation of stable hTERT-RPE1 cells expressing doxycycline-inducible UBE2H constructs (WT, AA and DD) by lentiviral transduction and application of CRISPR-Cas9 editing of the endogenous UBE2H locus to introduce the AA mutation (two independent clones, A3 and B4). **d.** Levels of doxycycline-inducible ubiquitin-charged UBE2H (WT, AA) were examined by anti-UBE2H immunoblotting in stably expressing hTERT-RPE1 cells. Cells were induced by increasing concentrations of doxycycline (0, 0.5, 1, 2.5, 5 and 10ng/mL). s.e., short exposure; l.e., long exposure. Quantification of the percentage of ubiquitin-charged UBE2H is shown (bottom). **e.** Levels of ubiquitin-charged endogenous UBE2H in WT cells and knock-in clones. Cells were treated with okadaic acid (200 nM) for 3h. Cell lysates were analyzed under non-reducing conditions to examine levels of UBE2H~Ub (top). Quantification represents mean ± s.d. from three independent biological replicates (bottom). **f.** Relative UBE2H mRNA levels in WT cells and knock-in clones. Quantification represents mean ± s.d. from three independent biological replicates. **g.** UBE2H protein levels in WT cells and knock-in clones with or without the proteasome inhibitor MG-262. Quantification of relative UBE2H protein level represents mean ± s.d. from three independent biological replicates. **h.** UBE2H degradation in WT cells and knock-in clones in the presence of cycloheximide (CHX) as measured by immunoblotting. Quantification of relative UBE2H protein level represents mean ± s.d. from three independent biological replicates. Statistical significance of differences between groups was determined with one-way ANOVA analysis (**f**) or two-tailed unpaired Student's t-test (**a, e, g-h**). *** = $p < 0.001$. ** = $p < 0.01$. * = $p < 0.05$. ns = not statistically significant.

### Extended Data Figure 3

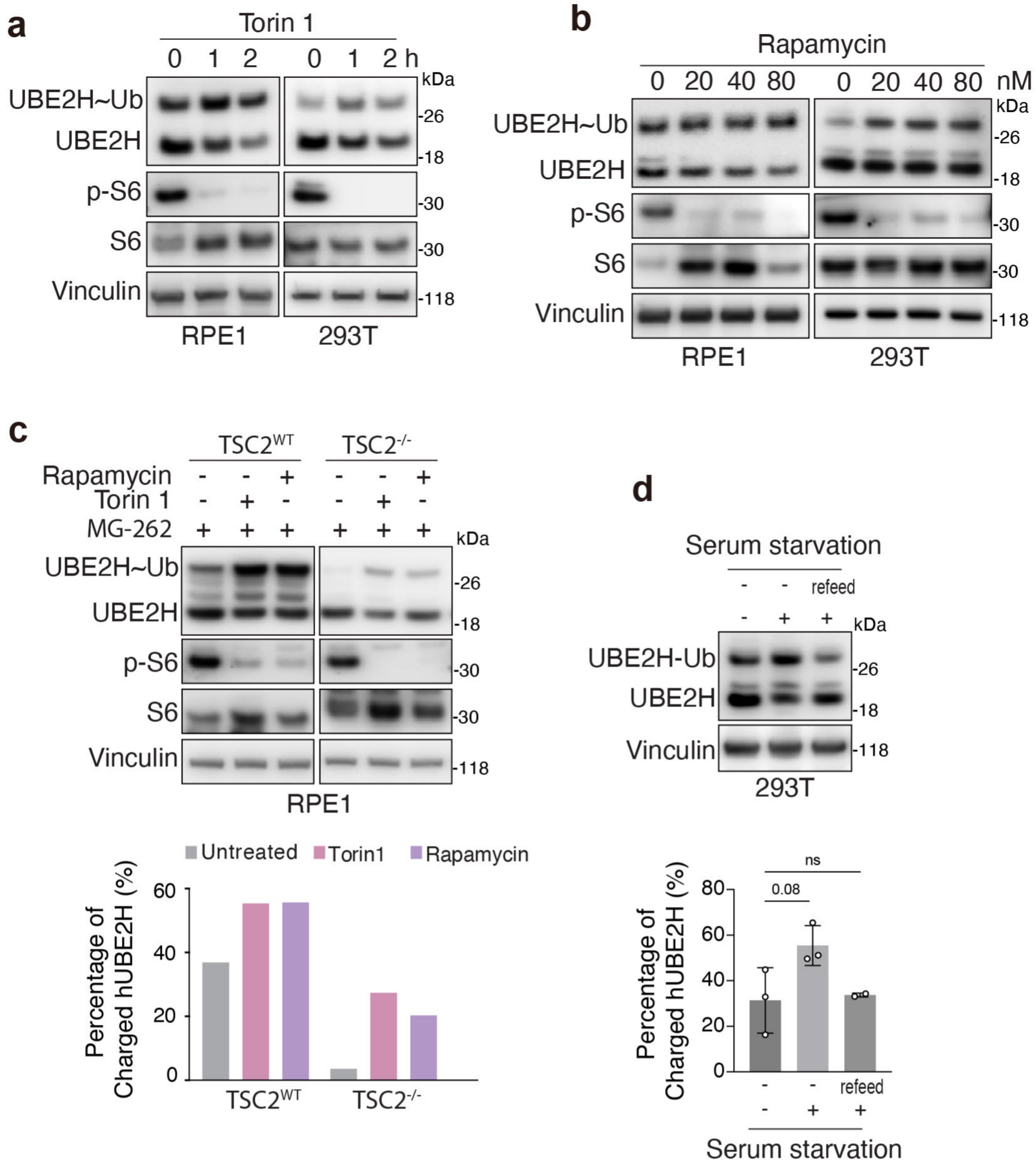


**Extended Data Fig. 3. a.** Non-reducing SDS-PAGE analysis of UBE2H thioester-linked ubiquitin conjugates in hTERT-RPE1and HEK293T cells after treatment with torin1 (250 nM) for the indicated time. Quantification represents mean ± s.d. from three independent biological replicates. **b.** Non-reducing SDS-PAGE analysis of UBE2H thioester-linked ubiquitin conjugates in hTERT-RPE1 cells and HEK293T cells after treatment with rapamycin at the indicated concentrations for 1 h. Quantification of the percentage of ubiquitin-charged UBE2H is shown (bottom). **c.** Non-reducing SDS-PAGE analysis of UBE2H thioester-linked ubiquitin conjugates after treatment with torin1 (250 nM) and rapamycin (80 nM) in WT, $TSC2^{-/-}$ hTERT-RPE1cells. Quantification of the percentage of charged UBE2H is shown (bottom). **d.** Non-reducing SDS–PAGE analysis of UBE2H thioester-linked ubiquitin conjugates in HEK293T cells that were incubated in serum-depleted medium for 12 h. Quantification represents mean ± s.d. from three independent biological replicates. Statistical significance of differences between groups was determined with one-way ANOVA analysis. * = $p < 0.05$. ns = not statistically significant.

### Extended Data Figure 4

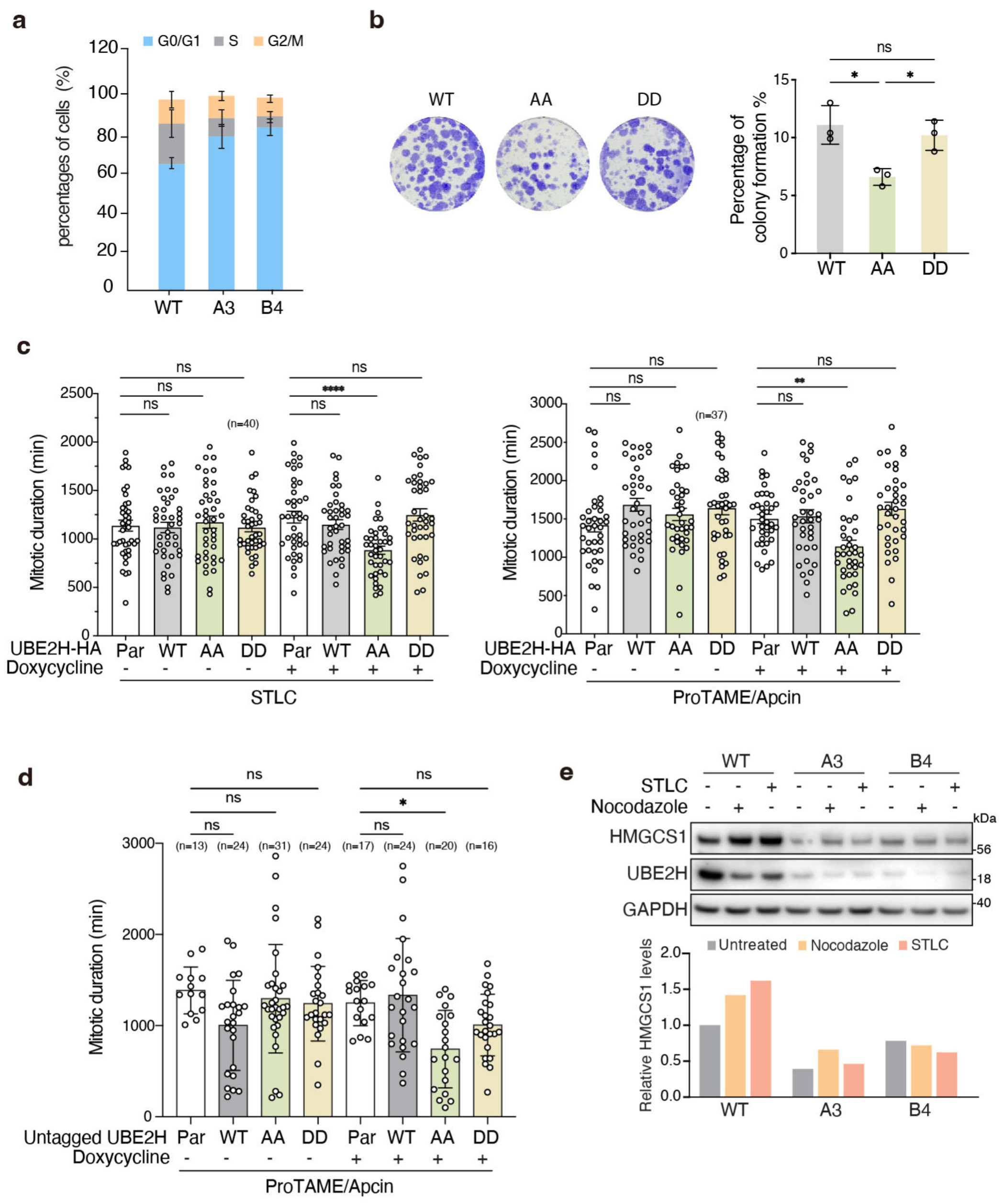


**Extended Data Fig. 4. a.** FACS analysis of propidium iodide (PI)-stained WT cells and knock-in clones (A3 and B4). Percentages of cells in each cell cycle phase were calculated. Quantification represents mean ± s.d. from three independent biological replicates. **b.** Colony formation assays in the hTERT-RPE1 cells that stably expressing doxycycline-inducible UBE2H constructs (WT, AA and DD) treated with doxycycline (5 ng/mL) for 14 days. Quantification represents mean ± s.d. from three independent biological replicates. **c.** Mitotic arrest duration of hTERT-RPE1 cells that stably express doxycycline-inducible UBE2H constructs (WT, AA and DD). Cells were treated with STLC (1.5 µM) or proTAME (12.5 µM) and apcin (25 µM) and imaged every 10 mins by wide-field time-lapse microscopy for 48 h. Quantification was performed on the number of cells indicated on the plot. Each point represents an individual cell's mitotic duration, measured as the time from NEB to division, slippage, or cell death. **d.** Mitotic arrest duration of hTERT-RPE1 cells that stably express doxycycline-inducible untagged UBE2H constructs (WT, AA and DD). Cells were treated with proTAME (12.5 µM) and apcin (25 µM) and imaged every 10 mins by wide-field time-lapse microscopy for 48 h. Quantification was performed on the number of cells indicated on the plot. Each point represents an individual cell's mitotic duration, measured as the time from NEB to division, slippage, or cell death. **e.** Western blots showing HMGCS1, UBE2H protein levels in asynchronous WT cells and knock-in clones. Mitotic cells were treated with nocodazole (600 nM) or STLC (2 µM) for 18h and collected by shake-off. Quantification of the relative HMGCS1 protein levels is shown (bottom). For b-c, Error bars = mean ± s.d. p-values were calculated by one-way ANOVA. **** = $p < 0.0001$. ** = $p < 0.01$. * = $p < 0.05$. ns = not statistically significant.

### Extended Data Figure 5

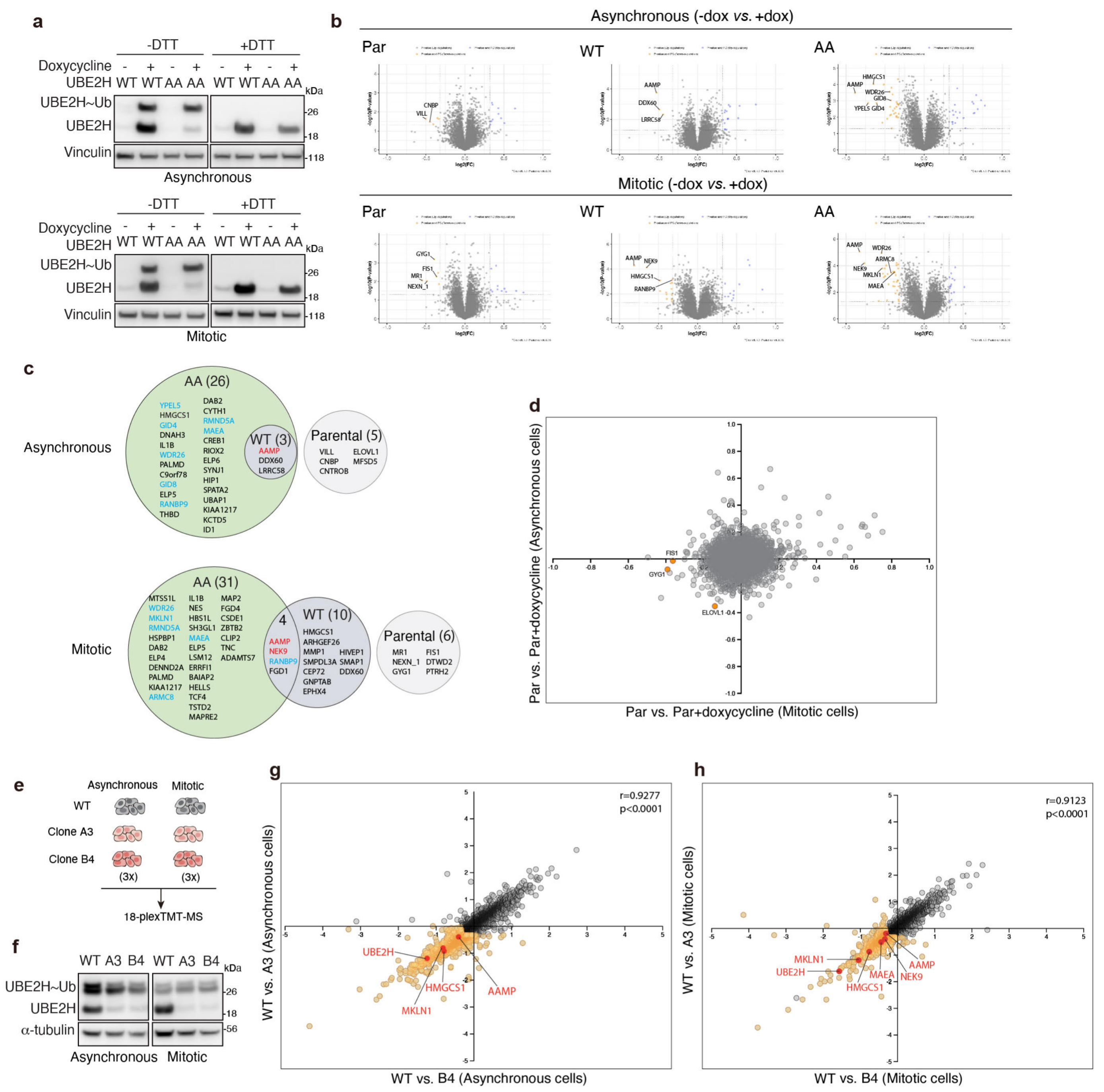


**Extended Data Fig. 5. a.** Western blotting of cells used for the proteomics experiment. Levels of doxycycline-inducible ubiquitin-charged UBE2H (WT, AA) were examined by anti-UBE2H immunoblotting in stable hTERT-RPE1 cells with or without the presence of DTT. **b.** Volcano plots of whole cell proteomic analysis of parental cells and UBE2H (WT, AA) cells treated with doxycycline compared with untreated cells in asynchronous and mitotic conditions. Shown are $-\log_{10}$ (P-value) (y-axis) vs $\text{Log}_2$ (fold change in protein abundance) (x-axis) for doxycycline-treated versus untreated cells, from three independent biological replicates. **c.** Venn diagrams showing overlap among proteins downregulated by doxycycline addition in parental cells or those overexpressing UBE2H (WT, AA) (FC < -1.25, p <0.05 and number of peptides >1), in asynchronous and mitotic conditions. **d.** Two-dimensional plots comparing proteins downregulated in parental cells after treatment with doxycycline in asynchronous and mitotic conditions. Significantly downregulated proteins are indicated in orange. **e.** Schematic of the TMT-mass spectrometry workflow to identify proteins differentially expressed in asynchronous and mitotic conditions in WT cells and knock-in clones across three independent replicates. **f.** Levels of ubiquitin-charged UBE2H were examined by anti-UBE2H immunoblotting in WT cells and knock-in clones in asynchronous and mitotic conditions. **g-h.** Two-dimensional plots of TMT-MS-based quantitation of protein abundance in the two knock-in clones under asynchronous and mitotic conditions. Each dot represents a quantified protein; axes indicate the log2-fold change in A3 (y-axis) and B4 (x-axis) relative to WT. Pearson's correlation analysis revealed a strong positive concordance between the two clones (asynchronous: r = 0.9277, p < 0.0001; mitotic: r = 0.9123, p < 0.0001). HMGCS1, MKLN1, UBE2H, AAMP, MAEA and NEK9 are highlighted in red, and the other downregulated proteins are indicated in orange.

### Extended Data Figure 6

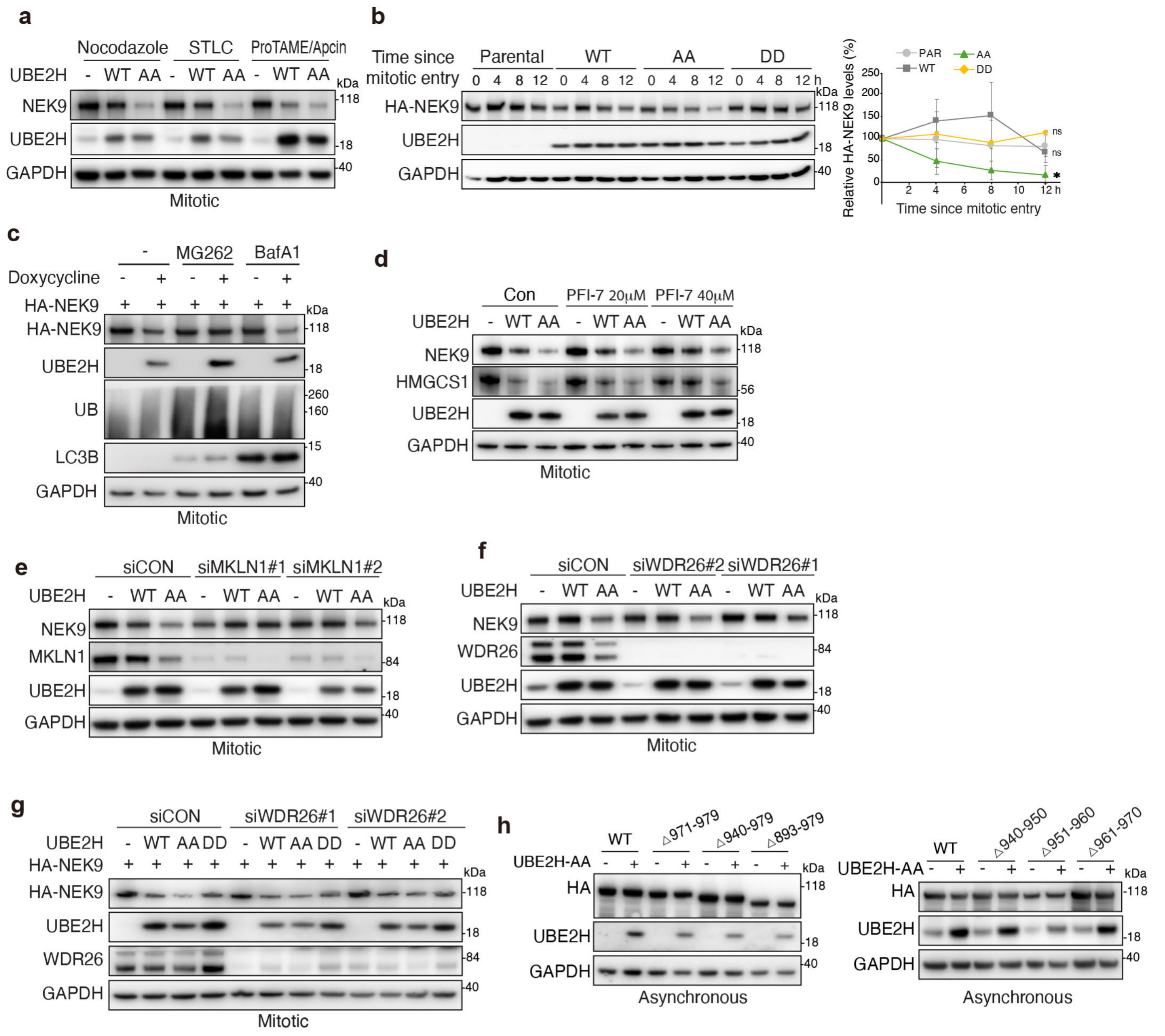


**Extended Data Fig. 6. a.** Western blots showing NEK9 protein levels in mitotic parental and UBE2H (WT, AA) hTERT-RPE1 cells after treatment with nocodazole (600 nM), STLC (2 μM), proTAME (12. 5 μM)/ apcin (25 μM). **b.** Immunoblots of NEK9 in hTERT-RPE1 cells. Cells were arrested by RO3306 7.5 μM for 18h, then released into nocodazole for the indicated time, and mitotic cells were harvested and analyzed by immunoblotting. Quantification represents mean ± s.d. from three independent biological replicates. **c.** Western blots showing exogenous NEK9 protein levels in mitotic UBE2H-AA-expressing hTERT-RPE1 cells after treatment with MG262 (10 μM) or BafA1 (1 μM) in the presence or absence of doxycycline. **d.** Western blots showing NEK9 protein levels in mitotic parental and UBE2H (WT, AA) hTERT-RPE1 cells after treatment with PFI-7 (40 μM). **e.** Western blots showing NEK9 protein levels in mitotic parental and UBE2H (WT, AA) hTERT-RPE1 cells transfected with siCON or MKLN1 siRNAs. **f.** Western blots showing NEK9 protein levels in mitotic parental and UBE2H (WT, AA) hTERT-RPE1 cells transfected with control or WDR26 siRNAs. **g.** Western blots showing exogenous NEK9 protein levels in mitotic parental and UBE2H (WT, AA, DD) hTERT-RPE1 cells transfected with siCON or WDR26 siRNAs. **h.** Western blots showing levels of NEK9 WT or truncation mutants in asynchronous UBE2H-AA-expressing hTERT-RPE1 cells in the presence or absence of doxycycline. Statistical significance of differences between groups was determined with two-tailed unpaired Student's t-test (b). * = $p < 0.05$. ns = not statistically significant.

### Extended Data Figure 7

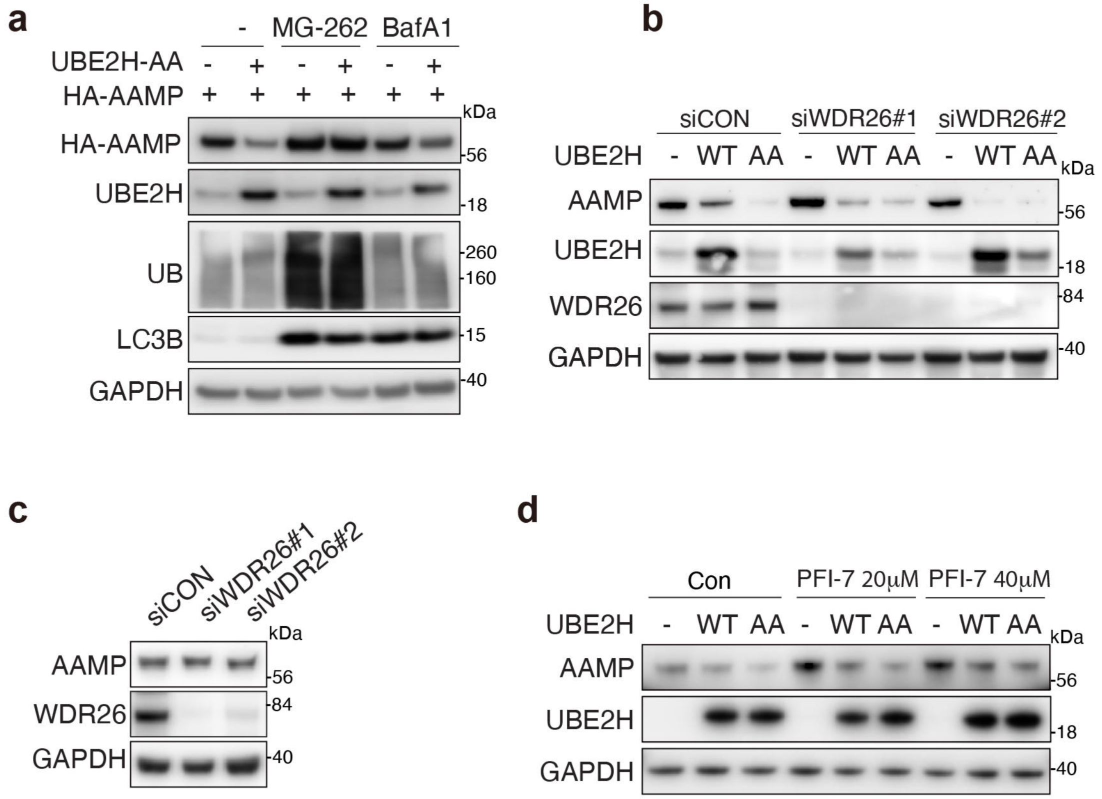


**e**

| | | |
|---|---|---|
| Human (Homo sapiens) | KVFCVQRPDR | 434 |
| Eastern newt (Notophthalmus viridescens) | KVFCLQRPDR | 417 |
| African clawed frog (Xenopus laevis) | KVFCVQQPDR | 443 |
| Cow (Bos taurus) | KVFCVQRPDR | 434 |
| Pig (Sus scrofa) | KVFCVQRPDR | 435 |
| Dog (Canis lupus familiaris) | KVFCVQRPDR | 435 |
| American crocodile (Crocodylus acutus) | KVFCVQRPDR | 436 |
| Mouse (Mus musculus) | KVFCVQRPDR | 435 |
| Rat (Rattus norvegicus) | KVFCVQRPDR | 435 |
| | *** (under PDR) | |

**Extended Data Fig. 7. a.** Western blots showing exogenous AAMP protein levels in UBE2H-AA-expressing hTERT-RPE1 cells after treatment with MG-262 (10 μM) or BafA1 (1 μM) in the presence or absence of doxycycline. **b.** Western blots showing AAMP protein levels in parental and UBE2H (WT, AA) hTERT-RPE1 cells transfected with siCON or WDR26 siRNAs. **c.** Western blots showing AAMP protein levels in hTERT-RPE1 cells transfected with siCON or WDR26 siRNAs. **d.** Western blots showing AAMP protein levels in parental and UBE2H (WT, AA) hTERT-RPE1 cells after treatment with PFI-7. **e.** Multiple sequence alignment of the indicated protein C-terminal region from representative vertebrate species. The conserved PDR motif is highlighted (boxed). Residue numbering corresponds to each sequence.

### Extended Data Figure 8

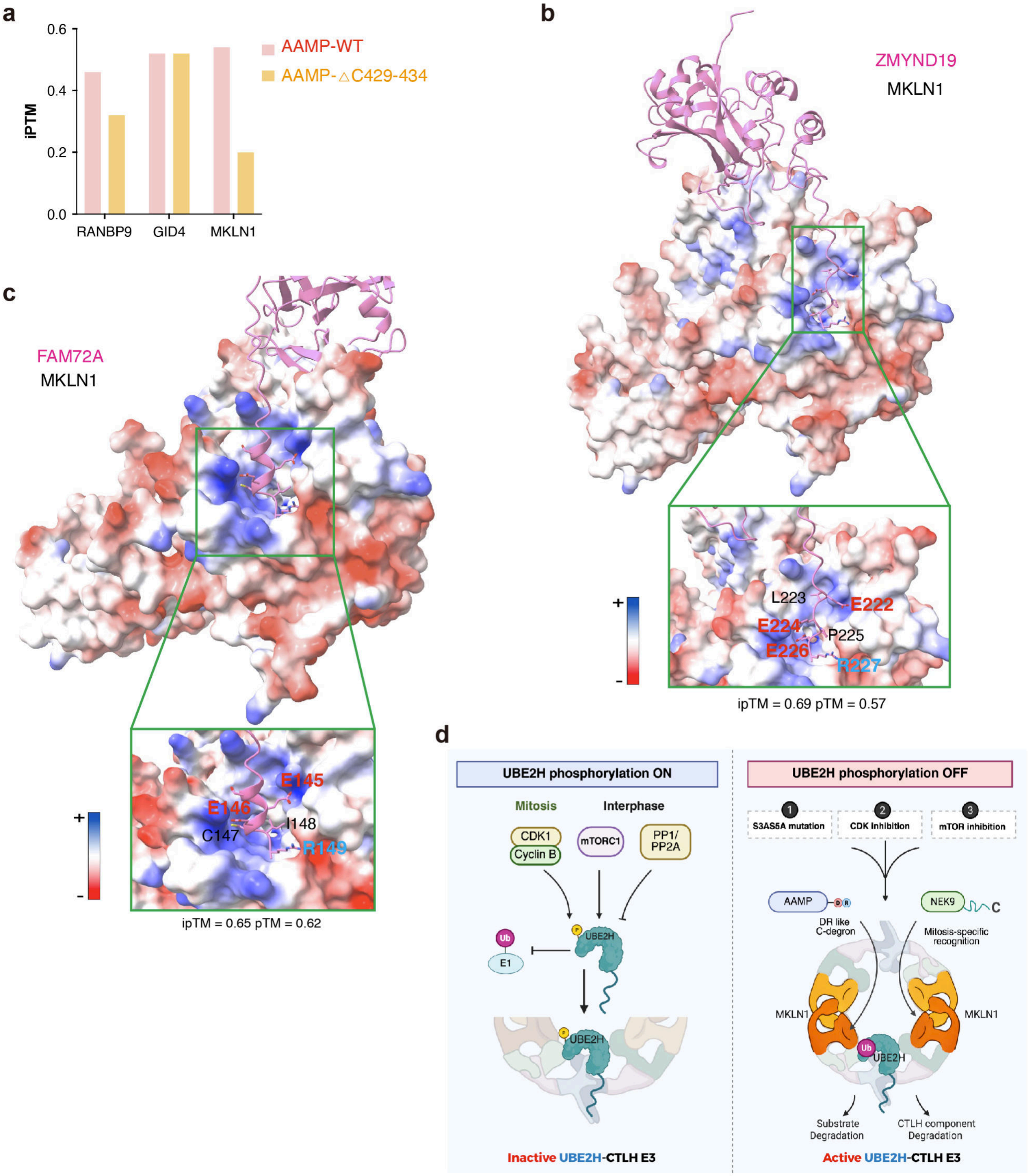


**Extended Data Fig. 8. a.** Predicted interactions between AAMP (WT, △C429-434) and CTLH components (RANBP9, GID4, MKLN1) using AlphaFold 3, showing ipTM scores. **b.** AF3 models for ZMYND19 (pink) bound to MKLN1, shown as an electrostatic potential surface. Residues at the extreme ZMYND19 C-terminus interact with a groove formed by the blades of MKLN1 β-propeller. ipTM scores for pairwise combinations are indicated. **c.** AF3 models for FAM72A (pink) and MKLN1 (electrostatic potential surface). Residues at the FAM72A C-terminus interact with the groove formed by blades of the MKLN1 β-propeller. ipTM scores for pairwise combinations are indicated. **d.** Graphical summary. Mitotic CDK- and interphase mTOR-dependent phosphorylation of UBE2H at N-terminal Ser3/Ser5 reduces UBE2H~Ub thioester formation (“charging”), thereby limiting the pool of active E2 available to CTLH. Preventing S3/S5 phosphorylation maintains UBE2H charging enhances CTLH-mediated substrate degradation (e.g. NEK9 and AAMP), and a DR-like C-terminal degron recognized by the CTLH subunit MKLN1 is defined.
